# Building better genome annotations across the tree of life

**DOI:** 10.1101/2024.04.12.589245

**Authors:** Adam H. Freedman, Timothy B. Sackton

## Abstract

Recent technological advances in long read DNA sequencing accompanied by dramatic reduction in costs have made the production of genome assemblies financially achievable and computationally feasible, such that genome assembly no longer represents the major hurdle to evolutionary analysis for most non-model organisms. Now, the more difficult challenge is to properly annotate a draft genome assembly once it has been constructed. The primary challenge to annotations is how to select from the myriad gene prediction tools that are currently available, determine what kinds of data are necessary to generate high quality annotations, and evaluate the quality of the annotation. To determine which methods perform the best and determine whether the inclusion of RNA-seq data is necessary to obtain a high-quality annotation, we generated annotations with 10 different methods for 21 different species spanning vertebrates, plants, and insects. We found that the RNA-seq assembler Stringtie and the annotation transfer method TOGA were consistently top performers across a variety of metrics including BUSCO recovery, CDS length, and false positive rate, with the exception that TOGA performed less in plants with larger genomes. RNA-seq alignment rate was best with RNA-seq assemblers. HMM-based methods such as BRAKER, MAKER, and multi-genome AUGUSTUS mostly underperformed relative to Stringtie and TOGA. In general, inclusion of RNA-seq data will lead to substantial improvements to genome annotations, and there may be cases where complementarity among methods may motivate combining annotations from multiple sources.

## INTRODUCTION

The reporting in 2001 of the first draft of the human genome sequence (Lander et al. 2001; Venter et al. 2001) ushered in a new era of genome-scale analysis, with a concomitant, rapid increase in the development of bioinformatics tools and resources to interrogate genomes for evolutionary patterns and features of biomedical interest. But even as genomes became available for other model organisms such as mouse (*Mus musculus*) (Mouse Genome Sequencing Consortium et al. 2002) and rhesus macaque (*Macaca mulatta*) (Rhesus Macaque Genome Sequencing and Analysis Consortium et al. 2007)—and had been previously published for smaller genomes such as *Drosophila melanogaster* (Adams et al. 2000)—the prohibitive cost of generating genome assemblies meant that research groups working on non-model organisms continued to operate in the genomic dark. Absent genome assemblies and annotations, such groups were forced to embark on time-consuming efforts to sequence small sets of conserved genes with Sanger sequencing using primers designed with other genomes, target anonymous loci such as AFLPs or *de novo* assembled RAD-seq reads. These methods imposed an analytical glass ceiling on the types of inferences that could be made and prevented the framing of research findings in a genomic context. While the advent of RNA-seq inched non-model organism research closer to understanding patterns at functional loci, *de novo* assembled transcriptomes presented novel analytical challenges and potential distortions of evolutionary patterns relative to what would be obtained with access to a genome assembly (Freedman et al. 2021).

Recent technological advances in long read DNA sequencing such as Pacific Biosciences HiFi (Wenger et al. 2019) and Oxford Nanopore (Jain et al. 2018), accompanied by dramatic reduction in costs have made the production of genome assemblies financially achievable and computationally feasible, such that genome assembly no longer represents the major hurdle to evolutionary analysis for most non-model organisms. Now, the more difficult challenge is to properly annotate a draft genome assembly once it has been constructed. The challenge is not so much the difficulty or computational resources required to run any one genome annotation tool, but how to a) select from the myriad gene prediction tools that are currently available, b) determine what kinds of data are necessary to generate high quality annotations, and c) evaluate the quality of the predicted transcript and gene models.

Currently available genome annotation tools approach the genome annotation problem in very different ways. Early computational tools for annotation used Hidden Markov Models (HMMs) to scan genomes for sequences representing protein-coding intervals, with AUGUSTUS (Stanke and Waack 2003) being the most widely used example. Recent implementations of this approach, such as BRAKER1 (Hoff et al. 2016) and BRAKER2 (Brůna et al. 2021) wrap optimized implementations of AUGUSTUS, using protein and RNA-seq evidence, respectively—and with the latest release, both—to train HMMs. Transcript assemblers such as Stringtie (Pertea et al. 2015) implement a graph-based framework to directly assemble transcripts from splice-aware alignments of RNA-seq reads to the genome. Tools such as Comparative AUGUSTUS (CGP) (Nachtweide and Stanke 2019) and TOGA (Kirilenko et al. 2023) use whole genome alignments to transfer annotation evidence between genomes, with the former involving multi-way transfer of HMM-based gene predictions, and the latter lifting over annotations from a high quality reference annotation in an exon-aware fashion.

A large part of difficulty in determining what strategy will work for “my genome” is annotation methods have, for the most part, been benchmarked and optimized with genomes from a very small slice of the tree of life: small genomes such as *D. melanogaster*, *Caenorhabditis elegans*, *Saccharomyces cerevisiae*, or with emphasis on vertebrates (Pertea et al. 2015; Hoff et al. 2016; Cantarel et al. 2008; Shao and Kingsford 2017) —no surprise given the implications for discoveries relevant to human health and disease. A related problem is that genome annotation will in the not too far future and, by necessity, need to be conducted by the group that has assembled a genome, rather than relying on Ensembl or NCBI to implement their automated pipelines. While the number of genomes annotated per year by NCBI has remained flat over the last few years, the number of published genome assemblies continues to increase at an unprecedented rate. In summary, publicly accessible generators of and repositories for annotations cannot keep up, such that the wait times between genome submission date and annotation completion will increase to the point of being impractical.

Motivated by the practical challenges facing research groups seeking to annotate new genome assemblies, here we evaluate the contents of genome annotations produced by ten different methods across a broad swath of the tree of life. These methods sample the current state of the art for different approaches to the annotation problem. Our taxonomic sampling includes two vertebrate clades (birds and mammals), two insect clades (*drosophila* spp., and heliconiine butterflies), and two plant clades (rosids and monocots), totaling 21 species. While “genome annotation” is often treated as an omnibus term that includes both the prediction of the genomic positions of genes and constituent isoforms, and the assigning of gene symbols and functions, we focus on the first of these two components. Our goals are to determine a) which methods are consistently top performers with respect to various sensitivity and specificity metrics b) the contents of individual annotations with respect to gene model fragmentation and fusion, c) whether the inclusion of RNA-seq data is essential for producing a high-quality annotation, d) whether species and taxonomic group affect annotation method performance, and e) whether there is complementarity among methods, such that integration of > 1 method might lead to detectable improvements in sensitivity.

## RESULTS

We evaluated ten different methods for genome annotation. At the core of five of these— MAKER (Cantarel et al. 2008) with both protein and RNA-seq evidence; BRAKER1; BRAKER2; CGP using protein evidence; and CGP with RNA-seq evidence—are HMM-based *ab initio* predictions by AUGUSTUS, with MAKER and BRAKER including predictions from additional *ab initio* tools. CGP enables evidence-based prediction within a species, as well as annotation transfer across species in the in the whole-genome alignment. TOGA performs an exon-aware liftover of annotations from a related genome that, ideally, has a complete, high quality genome annotation. Four methods assembly transcripts directly from RNA-seq alignments to a genome with one of two aligners: Stringtie with either HISAT2 (Kim et al. 2019) or STAR (Dobin et al. 2013) as aligner, and Scallop (Shao and Kingsford 2017) with either of these aligners. Many of our performance metrics use NCBI annotations as benchmarks. Derived from multiple lines of evidence, outputs of the NCBI annotation pipeline are good approximations to complete annotations. Futhermore, the species we annotated are either well-established model organisms or others with a long history of study and substantial genomic resources, such that their NCBI annotations are of the highest quality.

### BUSCO recovery

The ability of a gene prediction method to recover genes known to be conserved across a wide array of taxa is a good approximation for its ability to recover at least some sub-sequence of any gene, including those showing less conservation. BUSCO (Simão et al. 2015) scores – measuring the proportion of BUSCO targets in a search that have matches to query transcripts—varied considerably across methods and among species and broader taxonomic groups. Methods built on HMMs (BRAKER and CGP) consistently produced BUSCO scores for dipterans, heliconiine butterflies, and rosid plants that were higher than for other methods, or tied with TOGA (Fig. 1). In contrast, RNA-seq assemblers consistently outperform HMM-based methods in mammals and monocots, and one of the three bird species (Fig. 1). BRAKER_RNA-seq_ recovered more BUSCOs than BRAKER_protein_ in 16 of 20 species (Fig. 1), and in cases where

**Figure 1.**
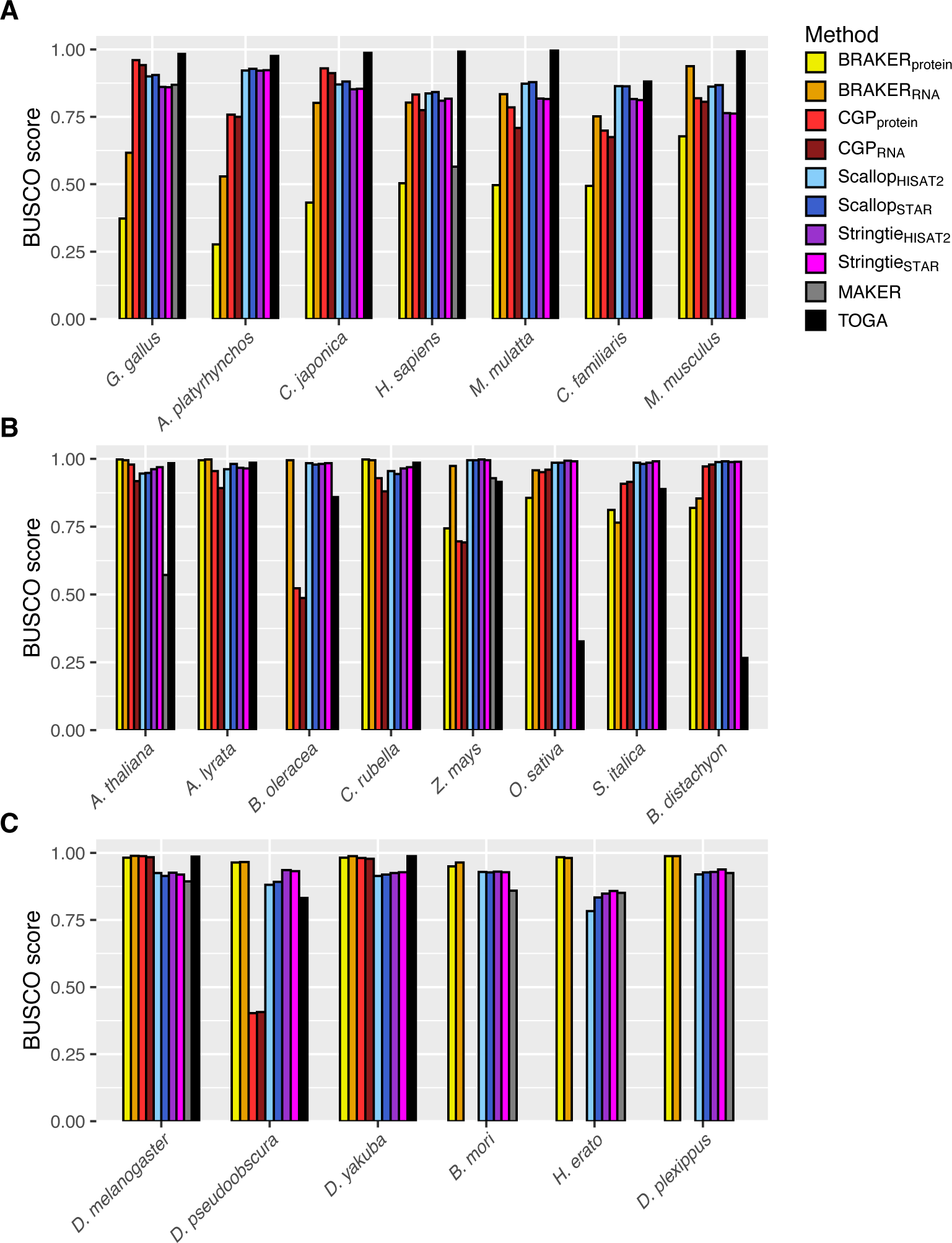
BUSCO scores by annotation method for (A) vertebrates, (B) plants,, and (C) insects.

BRAKER_protein_ recovered more the difference in recovery is typically small. BUSCO scores were very similar between CGP_protein_ and CGP_RNA-seq_, with CGP_protein_ recovering slightly more BUSCOs in 14 of 16 species (Fig. 1). CGP failed to produce more than a handful of transcript predictions for heliconiines, and BRAKER_protein_ consistently failed with *B. oleracea*, hence why these results are not included. For *D. pseudoobscura* and *B. oleracea*, CGP BUSCO recovery was poor, with BUSCO scores of approximately 50% or less. TOGA consistently produced the highest BUSCO scores in birds and mammals, and for most species in other groups had scores comparable to those produced by the top-performing method (Fig. 1). TOGA BUSCO scores relative to other methods was lower in plants, particularly in two of four monocots (Fig. 1). Somewhat surprisingly, MAKER, a putative full annotation workflow that leverages both protein and RNA-seq evidence, consistently lagged in performance behind other standalone methods. These patterns are not influenced by variation in the presence of BUSCOs in the underlying genome assemblies, which might produce taxonomic effects (correlation between BUSCO score and annotation BUSCO score/genome BUSCO score, Pearson’s ρ = 0.998, p = 2.2 × 10^−16^).

While there is considerable overlap between the BUSCOs that are recovered between classes of methods, there are species for which methods that use RNA-seq will recover BUSCOs that methods that use protein evidence cannot (Supplemental Fig. S1). This is particularly the case for species with larger genomes, such as birds, mammals, and *Z. mays*, the monocot in our sample with a genome > 2Gb, and that is six to 10-fold larger than the genomes of other monocots in our study. In some cases, 10-15% of BUSCOs are recovered by methods leveraging RNA-seq but not by methods relying on protein evidence. However, there are very few BUSCOs recovered by methods leveraging RNA-seq that TOGA does not also recover; the taxonomic exception is monocots, for which TOGA can perform poorly, likely due to known issues with whole genome alignment for plants.

### Annotation composition: number and length of CDS

The constituent CDS predictions that underlie BUSCO recovery rates varied widely among methods. Across diverse taxa, TOGA consistently produced CDS whose length distributions closely approximated those generated by NCBI, and with few exceptions Stringtie did as well (Supplemental Fig. S2). CGP annotations contained larger proportions of short predictions than other methods, with weaker trends towards shorter CDS observed in BRAKER. Scallop length distributions were often shifted towards shorter CDS to a similar degree as the HMM-based method with the greatest proportion of short CDS transcripts.

Annotations with tendencies towards shorter and far larger numbers of CDS relative to NCBI are likely indicative of fragmented transcript models. We observed this tendency for several species, particularly those with larger genomes (Fig. 2; Supplemental Fig. S3). Notable extreme outliers were the 7-fold and 4-fold larger CDS counts produced by CGP for macaque and human, respectively (Fig. 2A). The best approximations to median CDS length and number of NCBI annotations were typically produced by TOGA or a Stringtie assembly (Fig. 2; Supplemental Fig. S3).

**Figure 2.**
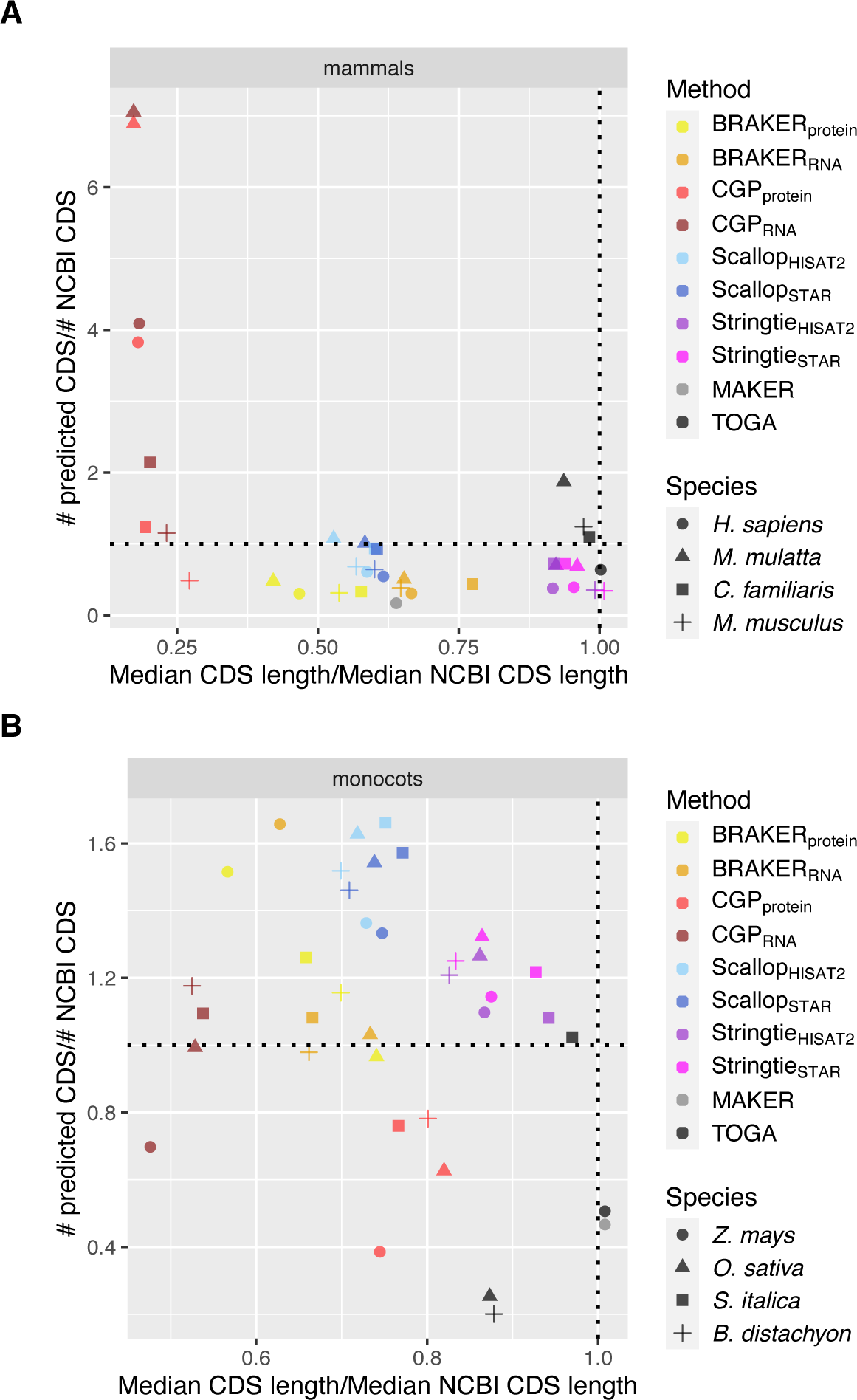
Joint distributions of number of predicted CDS (normalized by number of NCBI predictions) over median predicted CDS length (normalized by median NCBI CDS length) for (A) mammals, (B) monocots. Dotted lines indicated equivalence to NCBI annotation, such that methods that are closest to the intersection of those lines best approximate CDS length and number of NCBI annotations. For dipterans, birds and rosids, see Figure S3.

### False positives: intergenic predictions

For all but *A. thaliana*, the false positive rate (FPR) at which predicted genes fell entirely within intergenic regions relative to the respective NCBI annotations was lowest for TOGA; for *A. thaliana*, Stringtie (with STAR alignments) had the lowest FPR, and that for TOGA was nearly identical. In general, when TOGA did not have the lowest FPR, an RNA-seq assembler did (Fig. 3). For all species, FPR for the best performing method was ≤ 10% and for many species those predictions occurred < 5% of the time (Fig. 3). Regardless of the species and evidence type, FPR for CGP was much larger, often exceeding 50%, with predictions using RNA-seq evidence being less prone to FPR than those relying on protein evidence (Fig. 3). While BRAKER FPR was typically less than that of CGP, regardless of the evidence type used, FPR was higher than the best performing RNA-seq assembler in 16 of 18 species. The disparity between the low rates for RNA-seq assemblers compared to other methods was most evident in rosids, monocots, and mammals (Fig. 3). Nevertheless, the observed FPR suggest that, even for the best performing methods, hundreds or even thousands of gene predictions will fall outside of the genomic intervals for known real coding sequence. This raises the question of whether these sequences are false negatives in the NCBI annotation, or whether they are, in fact, false positives. We take a conservative approach and assume that most of these predictions are false positives, asking whether there are transcript features that might be predictive of such putative false positives so they can be removed. The same patterns held at the transcript level (gene vs. transcript intergenic prediction rate, Pearson’s σ = 0.994, p < 2.2 × 10^−16^).

**Figure 3.**
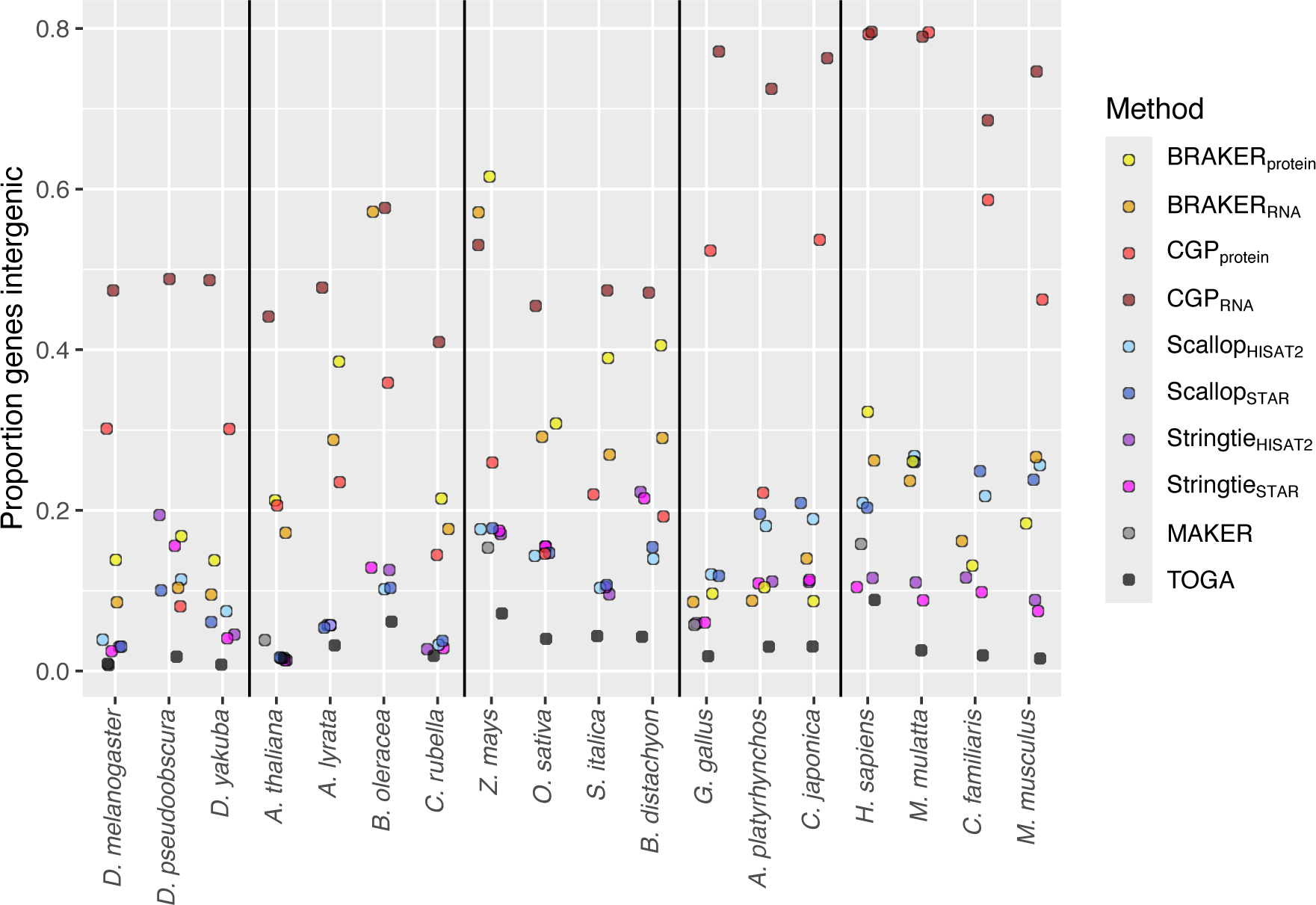
Proportion of predicted protein-coding genes that fall entirely outside the intervals for NCBI protein-coding genes, organized by species and method.

To better understand whether features of predicted transcripts distinguish those that at least partly overlap NCBI gene intervals from those that are entirely in NCBI intergenic regions, we fit random forest models to predictors summarizing sequence content, expression level, and whether the associated ORF had strong evidence to a match in the reference proteins of a related species. We did this for the five reference species for which NCBI annotations are thought to be the most complete. Overall, out-of-bag error rates (OOB) were consistently low for RNA-seq assemblers and TOGA, with OOB always being < 0.05 for TOGA, and for RNA-seq assemblers only exceeding 5% in *Z. mays* (Supplemental Fig. S4). Except for CGP, the class (genic vs. intergenic) error rates that comprise OOB are substantially higher for intergenic predictions (Supplemental Fig. S5). This result is undoubtedly due to the fact, that, for most species-method combinations, there are far fewer intergenic than genic predictions (Supplemental Fig. S6), making it harder for random forest to optimally classify intergenic sequences. However, random forest correctly classifies the majority of intergenic predictions as intergenic for methods that predict large numbers of intergenic transcripts (Supplemental Fig. S6).

Estimates of node purity by the Gini index—an estimate of variable importance that quantifies the extent to which removing a predictor reduces the frequency of predictions matching the true class on either side of a split—reveal that whether or not a transcript ORF has a BLAST hit is frequently the most powerful predictor of whether or not a transcript is genic or intergenic (Supplemental Fig. S7), with expression level and CDS length also frequently making large contributions to the models. These results suggest that, in the absence of a truth-set of known CDS intervals, some CDS features are potentially useful for setting filters to discriminating true from false positive intergenic predictions. The importance of individual expression metrics is likely underestimated, due to the correlation between expression metrics, such that the effect of removing one is mitigated by the presence of another in any one of the constituent trees in the random forest. These results collectively suggest that short, lowly expressed transcripts without hits to an external protein database are enriched for intergenic (and likely spurious) predictions.

### Gene fusions

We defined fusions as cases where for a predicted gene, the CDS of an associated transcript overlapped with the CDS of > 1 NCBI gene, or different CDS transcripts of a predicted gene each overlapped with the CDS of a single NCBI gene, but different predicted CDS transcripts overlapped with different NCBI genes; these definitions were not mutually exclusive. With few exceptions, the rate of putatively false fusions fell below 5% across species and methods, with notable exceptions for a handful of MAKER and BRAKER_RNA_ gene sets (Fig. 4). In general, fusion rates for HMM-based methods were lower than for RNA-seq assemblers, likely due to the tendency for shorter CDS lengths of the former; TOGA fusion rates were consistently among the lowest (Fig. 4). Gene-level fusions were not merely due to individual predicted CDS transcripts spanning the CDS of multiple NCBI genes, but were often dominated by cases where different transcripts from the same predicted gene each overlapped the CDS of a different NCBI gene (Supplemental Fig. S8).

**Figure 4.**
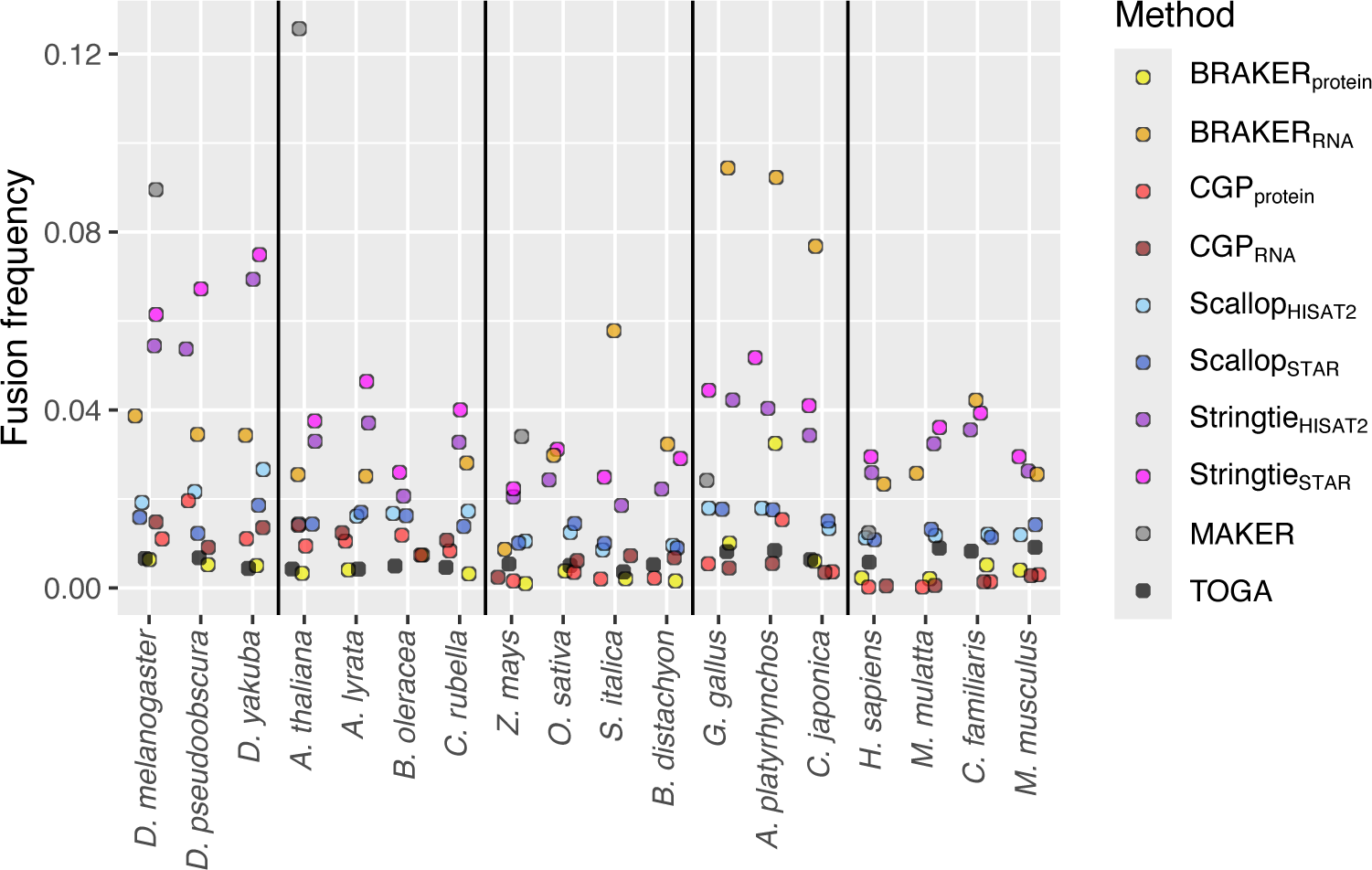
Frequency of gene fusions by species and method. Fusions are defined as when individual transcripts overlap the CDS of multiple NCBI genes, different transcripts from the same predicted gene each have overlaps to different genes, or a combination of both of these. Frequencies are calculated after filtering out NCBI-annotated fusion events.

### Protein sequence completeness

BRAKER and CGP consistently had the highest percentage of predicted proteins that had proper start and stop codon without any internal stop codons (Supplemental Fig. S9), with the former usually outperforming the latter by a modest margin. TOGA predictions had up to 20% fewer complete proteins than these methods, with Stringtie performing either slightly better or slightly worse, depending upon the species. Scallop consistently had the smallest percentage of predictions that represented complete proteins. Despite being a pipeline meant to process predictions from multiple *ab inito* tools (e.g. AUGUSTUS), MAKER performed worse than BRAKER and CGP and was often, and for plants had the smallest proportion of complete protein predictions. Nevertheless, most predicted transcripts in an annotation have a structure consistent with complete proteins, and with the exception of Scallop, rarely dipped below 80%.

The utility of “completeness” may be misleading as a stand-alone measure of whether a prediction is correct (or representing a true protein-coding sequence), if the wrong trinucleotide sequences are classified as start and stop codons due to incorrect inference of splice sites. That this may happen is highlighted by contrasting rates of completeness with the frequency with which predicted protein sequences match those in high quality protein databases. For each species, BLASTP was performed against a database comprised of proteins derived from the NCBI annotations of the species we included in their taxonomic group, including the species in question. The overall proportions of BLASTP hits were, with the exception of CGP, very high (Fig. 5A). TOGA was consistently the top performer, but for dipterans and rosids, TOGA, RNA-seq assemblers and BRAKER were barely distinguishable; for monocots, birds, and mammals,

**Figure 5.**
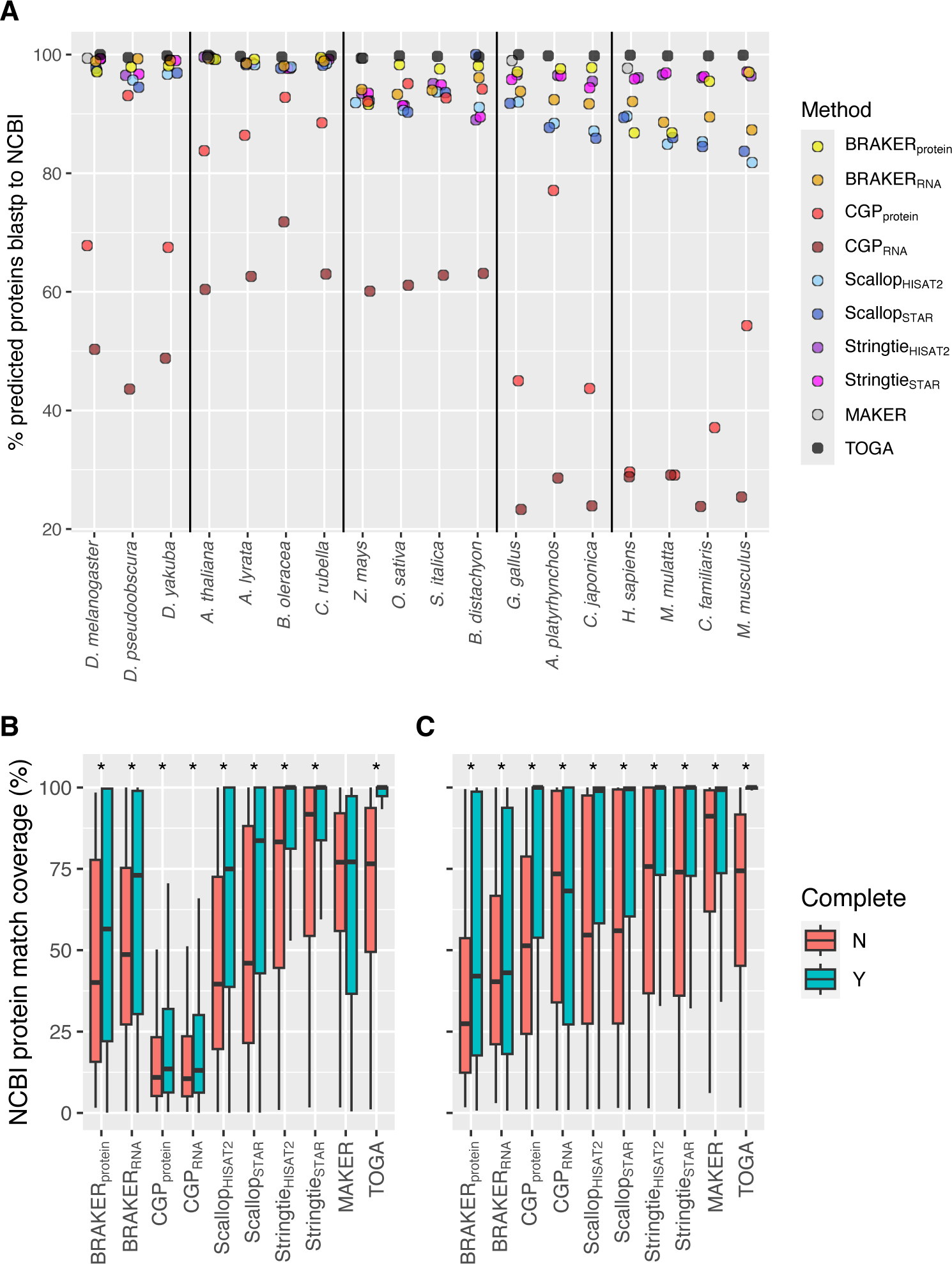
(A) By species and method, proportion of predicted proteins with BLASTP hit to NCBI proteins of species in taxonomic group. For predicted proteins with BLASTP hits, proportion of NCBI best hit target covered by amino acid matches with the predicted protein for (B) *H. sapiens* and (C) *Z. mays,* broken into proteins that are complete (start and stop codons present, no internal stop codons), and those that are not. Benjamini-Hochberg adjusted p-values for Wilcoxon rank-sum tests p≤0.05 indicated by *.

BRAKER with protein evidence was the second highest rate of BLASTP hits, with Stringtie following close behind (Fig. 5A). However, when we focused on our reference species for which NCBI annotations are of highest quality, in looking at the distributions of the coverage of NCBI proteins in BLASTP hits—defined as the number of matching bases by the length of the best-hit target—with the exception of *D. melanogaster*, there was broad overlap in coverage distributions between complete and incomplete protein predictions for BRAKER, CGP, and MAKER (Figs. 5B, 5C; Supplemental Fig. S10). This suggests these tools are frequently producing truncated protein predictions by identifying the wrong start or stop codons. In contrast, there was far less overlap for RNA-seq assemblers and TOGA. Stringtie and TOGA appear to do a better job than other tools of reconstructing the amino acid sequences of high-quality reference annotations that we defined as our truth set. The strong performance of TOGA cannot be attributed solely to the fact that the annotations being transferred originate from one of the species contained in our protein databases used for BLASTP searches, as excluding the proteins originating from the species whose annotations are being transferred led to negligible decreases in percentages of predicted proteins with hits: *H. sapiens*, 99.9% vs. 98.9%; *G. gallus*, 100% vs. 99.7%; *D. melanogaster*, 100% vs. 99.7%; *Z. mays*, 99.4% vs. 99.3%; and *A. thaliana*, 99.9% vs. 99.8%.

### Transcriptome representation: expression

Random forest models we constructed to distinguish intergenic predictions from ones overlapping known protein-coding gene intervals indicate that the former may be characterized by low expression. Predictions by BRAKER and CGP (regardless of the evidence type) and to a lesser extent MAKER contain larger proportions of genes with TPM < 1 (Fig. 6A; Supplemental Fig. S11) relative to RNA-seq assemblers. HMM-base methods using RNA-seq evidence often had larger or comparable fractions of lowly expressed genes compared to their counterparts using protein evidence (Fig. 6A; Supplemental Fig. S11). TOGA predictions did not have substantially elevated proportions of low TPM genes (relative to RNA-seq assemblers), and for monocots had the lowest fraction of lowly expressed genes (Fig. 6A; Supplemental Fig. S11). This may be due to alignments of expressed sequences being part of the evidence used by the NCBI annotation pipeline. In general, with their direct connection to expression, transcript assemblies of RNA-seq data (particularly those based upon Stringtie) consistently had the smallest fraction of lowly expressed transcripts.

**Figure 6.**
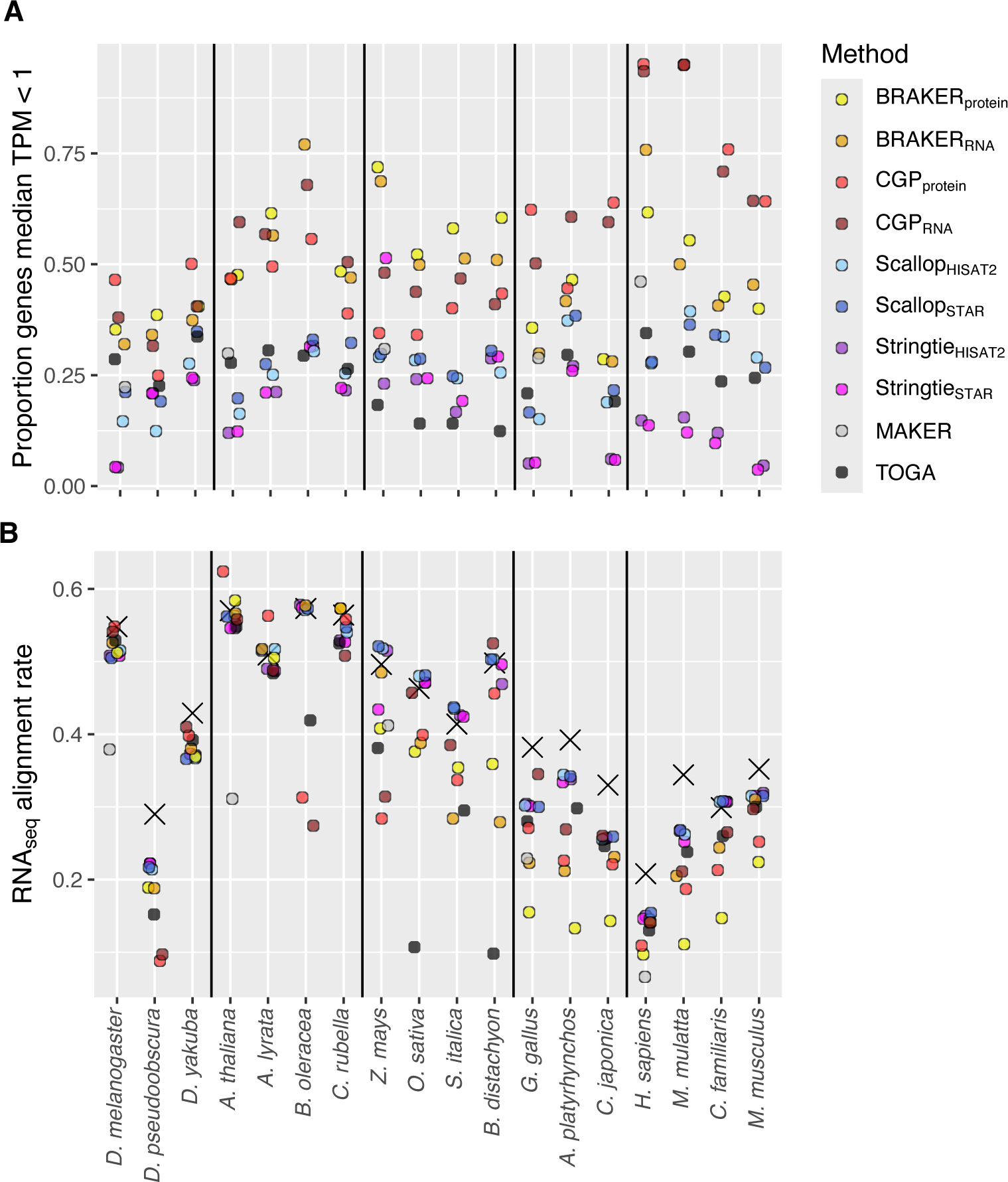
By species and method, (A) proportions of predicted genes for which TPM < 1, and (B) RNA-seq read alignment rates to predicted transcripts. For Stringtie and Scallop, UTR intervals have been removed, such that all annotations are for CDS only. Xs denote rates for the NCBI annotations (also excluding UTR intervals).

The RNA-seq read alignment rate to an annotation provides an estimate of the extent to which an annotation captures the underlying expressed transcriptome. Because BRAKER2, CGP, MAKER, and TOGA do not predict UTRs, and because BRAKER1’s UTR prediction option was an experimental features we chose not to include, we removed UTR intervals from Stringtie and Scallop annotations to make methods comparable. When UTRs were excluded from consideration, alignment rates were relatively low, only ever exceeding 50% for rosids, *D. melanogaster,* and for two of four monocots (Fig. 6B); this is partly due to the RNA-seq data containing reads originating from UTR intervals. While RNA-seq assemblers had alignment rates that were frequently the highest for a particular species, there were only modest differences between these tools and the best performing implementations of BRAKER or CGP (Fig. 6B). Rates for TOGA were lower than these, and were at approximately 10% for two of four monocots, with rates for MAKER also being low relative to most other tools (Fig. 6B). Regardless of method, alignment rates for human annotations always fell below 20%. Even so, alignment rates varied in a manner similar to NCBI annotations without UTRs (Pearson’s ρ = 0.86, p=2.2 × 10^−16^), albeit in most species-by-method combinations lower than those for NCBI (Fig. 6B).

Because these comparisons were based upon alignment of the same reads that were assembled by Stringtie and Scallop, we considered the possibility that this would provide an unfair advantage to the assemblers relative to tools that only used RNA-seq data to generate splice hints (BRAKER_RNA_ and CGP_RNA_), or to filter HMM-based predictions post hoc (MAKER), and even more so for methods that did not use RNA-seq data (BRAKER_protein_ and CGP_protein_). Training and test data alignment rates were strongly correlated (Pearson’s ρ = 0.86, p=2.2 × 10^−16^), although for all species except human (and for the BRAKER_protein_ annotation for chicken), test alignment rates were lower than training rates (Supplemental Fig. S12A). Nevertheless, the reduction in test data alignment rates relative to training data were modest, only exceeding 10% for *D. melanogaster* and *Z. mays* for several methods (Supplemental Fig. S12B). Only for *D. melanogaster* was there a clear drop in the test alignment rate for Stringtie and Scallop relative to other methods (Supplemental Fig. S12), suggesting that the recycling of training data for alignment does not generate substantial bias in the broad patterns we observe.

RNA-seq assemblers are agnostic to the functional role of the sequence intervals from which reads originate, while HMM-based approaches and TOGA do not predict UTRs. The inclusion of UTR intervals predicted by the assemblers led to large increases in their alignment rates, such that they outperformed other methods (Supplemental Fig. S13). The magnitude of this alignment rate difference raises questions regarding the contents of predicted UTR intervals. We thus assessed whether the lengths of predicted UTRs are consistent with those found in NCBI annotations, and whether the alignment rate increase they provide is due, at least in part, to there being true CDS intervals contained within UTRs predicted by Transdecoder— CDS that it failed to predict. We address these questions immediately below.

### UTRs in RNA-seq assemblers

Ratios of UTR to CDS length are greater for Scallop and Stringtie assemblies than for NCBI annotations (Supplemental Fig. S14). These disproportionately long (relative to NCBI) UTRs constitute an excess of target sequence for alignment, such that their exclusion clearly contributes to a large drop in alignment rates. We considered the possibility that the greater proportional length of UTRs for RNA-seq assemblers relative to NCBI annotations could be due to the NCBI pipeline computationally truncating UTRs when mitigating for cases of putative transcriptional readthrough past stop codons. In this case the gold standard genomes (*H. sapiens*, *M. musculus*, *D. melanogaster*, *Z. mays*, and *A. thaliana*) for which there has been extensive curation would be expected to have a lower ratio of UTR to CDS length. However, plotting the ratio of predicted UTR to CDS ratios for the assemblers over that for NBCI predictions produces the opposite pattern, where these ratios of ratios are lower for the gold standard genomes (Supplemental Fig. S14).

Next, we evaluated whether Transdecoder pipeline fails to predict CDS at the ends of transcripts, and be default assigns them to the UTR functional class. If this were a pervasive problem, then we would expect a large fraction of UTRs to have BLASTX hits to an NCBI protein database for the same species. As would be expected if undetected CDS occur in the UTR intervals for RNA-seq assemblers, the percentage of transcripts with a UTR BLASTX hit as the length of UTRs relative to CDS for assemblers increases relative to that observed for NCBI annotations (Fig. 7). Depending upon the species and particular assembler-aligner combination, up to 60% of transcripts may potentially contain undetected CDS in regions annotated as UTRs. This suggests the failure to detect CDS plays a substantial role in the drop in RNA-seq alignment rates when UTRs are excluded. This then leads to an underestimate of the realizable alignment rate for RNA-seq assemblers than if these CDS were properly incorporated into the annotations. It seems likely that the frequency of such undetected CDS exons may contribute to the tendency for assemblers to have lower percentages of proteins classified as complete relative to HMM-based methods (Supplemental Fig. S9).

**Figure 7.**
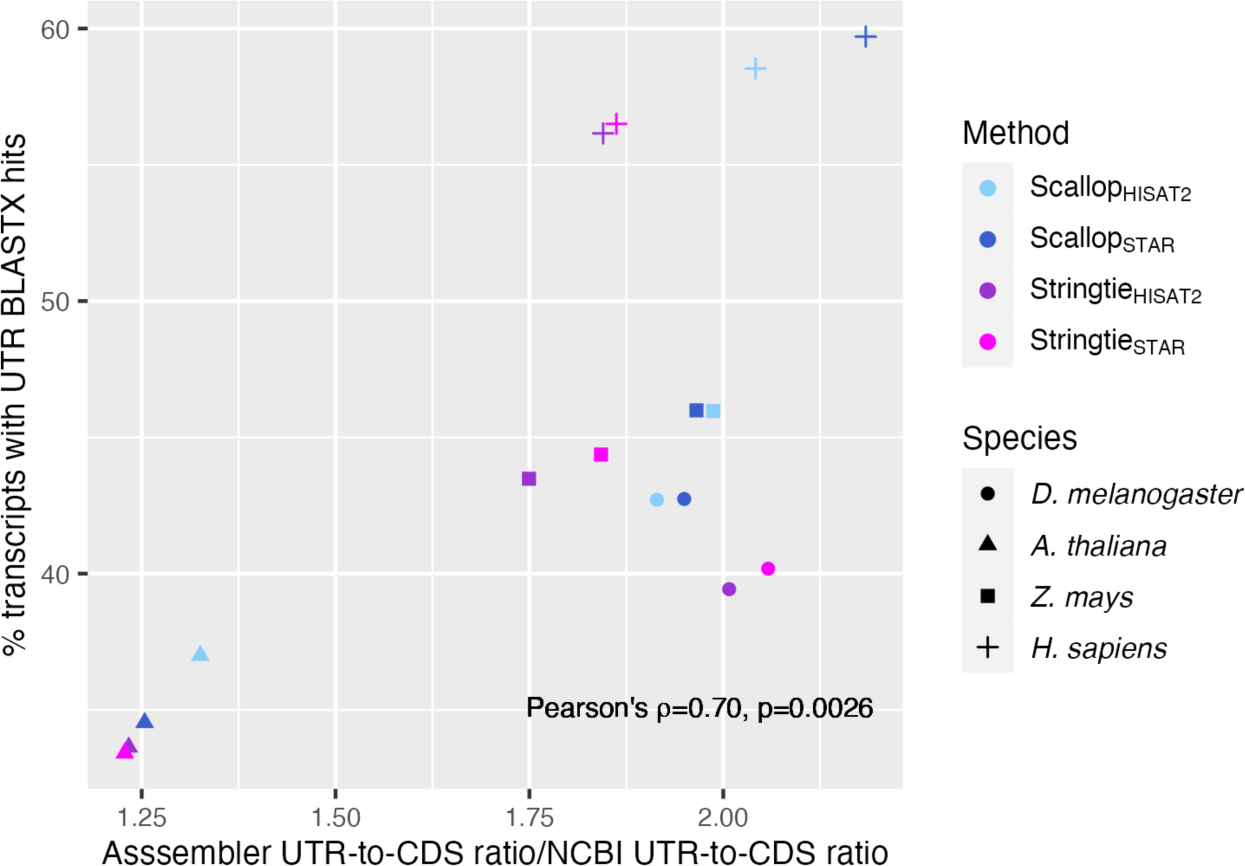
Evidence for undetected CDS in predicted UTR intervals for Stringtie and Scallop. Increasing percentage of RNA-seq assembler transcripts with UTRs that have a BLASTX hit to the NCBI protein database of the same species, as a function of an increase in the assembler UTR-to-CDS ratio relative to that for NCBI annotations.

### Multi-method integration

Above, we have demonstrated that MAKER, a workflow for integrating and filtering predictions from multiple *ab initio* prediction tools, underperforms compared to several different stand-alone annotation methods and across a variety of metrics. Another integration approach, TSEBRA (Gabriel et al. 2021a) selects transcripts and merges transcript models from separate runs with protein and RNA-seq evidence, respectively. TSEBRA integration produces an annotation with BUSCO scores that are no better than the best of the two BRAKER runs (Supplemental Fig. S15A), and in most cases have slightly worse scores. Proportions of predicted transcripts with BLASTP hits to the NCBI proteins from the species group we investigated are consistently worse than either BRAKER implementation (Supplemental Fig. S15B). Similarly, in four of five reference species, the percentage of intergenic genes is higher for TSEBRA than either BRAKER implementation (Supplemental Fig. S15C), with RNA-seq alignment rates falling between the two BRAKER runs (Supplemental Fig. S15D). In short, MAKER and TSEBRA lead to a loss of meaningful transcriptome information relative to the best of the alternative BRAKER approaches.

While not necessarily an optimized method for merging annotations, we investigated whether adding genes (and their constituent child features) from a second annotation to a base annotation—requiring that those added genes fell entirely outside of the gene intervals of that base annotation—would leverage the complementary strengths of different methods while minimizing information loss. We also explored whether successive additions from different methods would continue to improve the overall annotation. There was a high degree of variability in the benefits of such integration. While stand-alone methods like TOGA had the highest BUSCO score, adding TOGA annotations to those of Stringtie achieved the best balance of maximizing both BUSCO score and RNA_seq_ alignment rate, overcoming the tradeoff observed between these two metrics often seen in individual methods (Fig. 8A). Which method one chooses as the base annotation can impact BUSCO scores, the number of genes that have BLASTP hits to NCBI proteins, the percentage of predicted genes with such hits, and the RNA-seq alignment rate. For example, using Stringtie_STAR_ as a base annotation leads to lower BUSCO scores and higher alignment rates than for integrations that start with TOGA (Figs. 8A, 8B; Supplemental Figs. S16, S17), and while integration increased, as expected by definition, an increase in predicted genes, the percentage of genes with at least one transcript having a BLASTP hit to an NCBI protein decreased, suggesting that both real and spurious predictions get added (Supplemental Figs. S16-S19). Method integrations that did not include TOGA produced high RNA_seq_ alignment rates but lower BUSCO scores than TOGA and integrations that used it as the base annotation (upon which to add others), with reasonably high fractions of genes with BLASTP hits (Fig. 8C; Supplemental Fig. S18). Finally, in a scenario where RNA-seq data is not available, adding BRAKER_protein_ annotations to those of TOGA did little else other than to minimally increase the RNA_seq_ alignment rate (Fig. 8D; Supplemental Fig. S19). In these four scenarios, adding BRAKER_protein_ had negligible effect.

**Figure 8.**
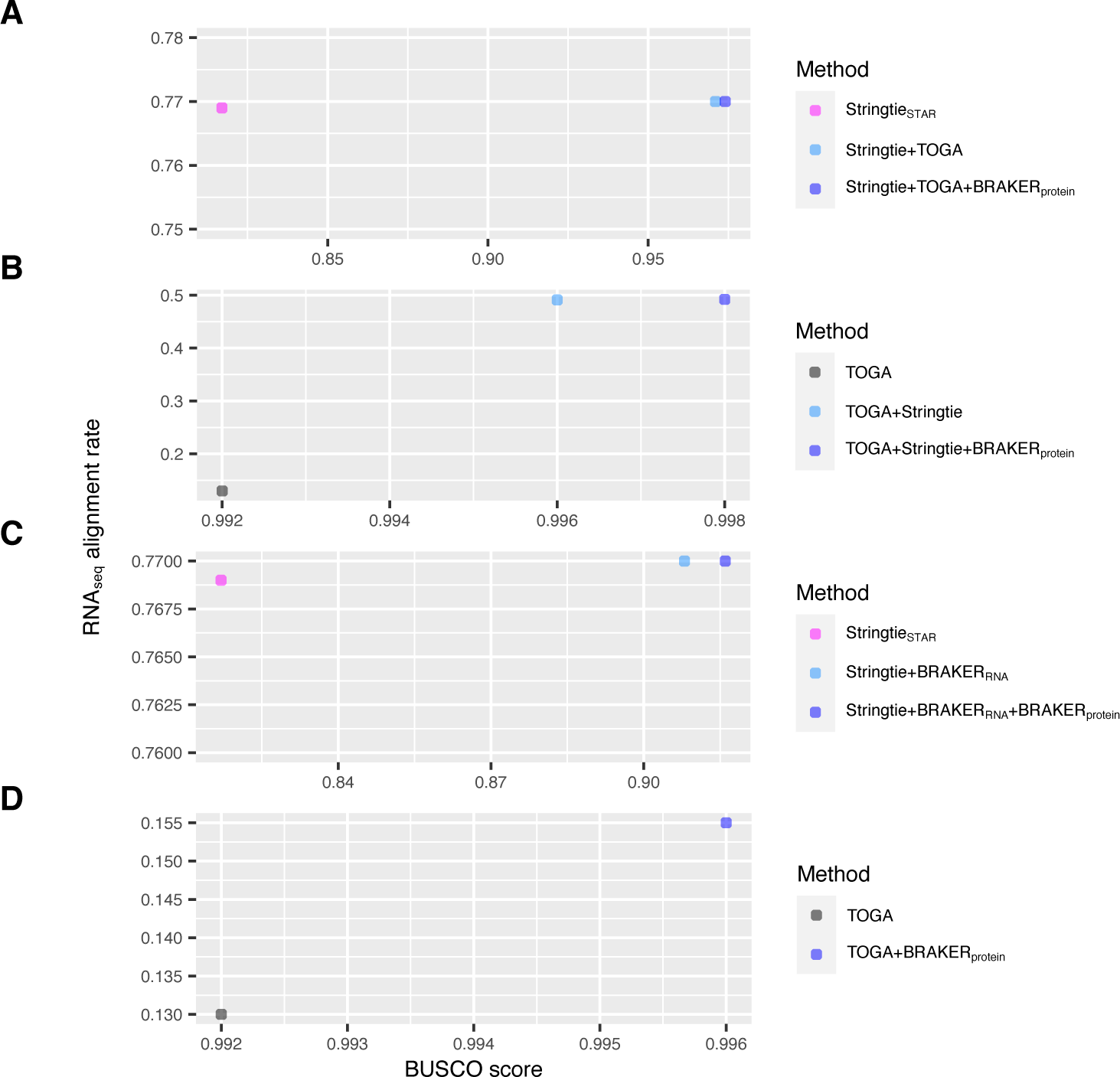
Bi-plots of median sample RNA-seq alignment rate versus BUSCO score for individual base annotation methods and subsequent integrations for (A) Stringtie_STAR_ as the base, with subsequent integration of TOGA and BRAKER_protein_ (B) same as B but with TOGA as the base annotation, (C) the absence of a closely related high quality genome annotation, precluding the use of TOGA, with Stringtie_STAR_ set as the base annotation, and (D) no RNA-seq data, with TOGA as the base annotation, and an integration of BRAKER_protein_. Integration involves successively adding genes from one annotation that fall entirely outside of the annotation to which those genes are being added.

## DISCUSSION

With genome assembly and annotation increasingly becoming part of the workflow for researchers studying non-model organisms, the choice of an annotation method depends upon knowing whether a particular method will perform well in the species in question, as well as what data will need to be generated to generate the best quality annotation possible. Previous efforts to evaluate and compare methods have sampled a small slice of the tree of life, and often focused on small, tractable genomes with extensive genomic resources, or other model organisms such as *H. sapiens* or *M. musculus*. This can make choosing a method more difficult, because the species used to benchmark annotation tools may be evolutionary distant from a newly assembled genome of interest. Our investigation overcomes this problem by evaluating a large set of methods across the broadest taxonomic swath investigated to date, revealing both cross-species and taxon-restricted patterns. By identifying methods that performed well across diverse species, we believe that researchers using those methods will likely be able to generate reasonably high-quality annotations for their newly assembled genome of interest. Our findings have implications for study design, data collection, and annotation method choice, and highlight ongoing challenges that require further methods development.

First and foremost, the inclusion of RNA-seq data will invariably improve annotation quality, particularly when used to assemble transcripts directly from sequence alignment with Stringtie. While HMM-based methods such as CGP and BRAKER using either protein or RNA-seq evidence can produce high BUSCO scores, they consistently lag behind RNA-seq assemblers with respect to their representation of the underlying transcriptome—as characterized by read alignment rates. Furthermore, HMM-based approaches typically produce thousands of false-positive predictions which need to be filtered but may be difficult to identify. That HMM-based predictions may consistently be “complete”, with proper start and stop codons, yet also shorter than NCBI transcripts and those produced by RNA-seq assemblers, highlights a shortcoming of HMM-based methods—splicing patterns that are consistent with an inferred model of protein-coding sequence structure may not be the real pattern, instead representing truncated open reading frames or spurious predictions. While we did not assess the utility of cDNA long reads for annotation, we expect our findings to be robust to their adoption, and that the performance disparities between RNA-seq assemblers and HMM-based methods will almost certainly widen. The direct evidence of splicing patterns across full length reads will enable reconstruction of full-length transcripts, while in the HMM context, longer reads will simply lead to more accurate detection of splice sites, and more accurate model parameterization—to the extent that any model can capture the diversity of sequence composition and splicing patterns observed in higher organisms. We suspect that increasingly accurate model parameterization will lead to diminishing returns relative to direct assembly of transcripts from reads. RNA-seq assemblers are particularly valuable for larger, more complex genomes and consistently rank among the top methods across diverse performance metrics. Furthermore, and while not the focus of this study, assemblers permit the inclusion of non-coding RNAs in an annotation, with the caveat that it is more difficult to distinguish real non-coding transcripts from spurious assembly of low-coverage transcriptional noise. Overall, while the advantages of using RNA-seq assembly over other methods may be diminished for smaller, less complex genomes, it clearly produces better annotations than HMM-based approaches for more complex genomes, and reliably produces relatively complete annotations regardless of taxonomic group or genome organization.

The demonstrated superiority of RNA-seq assemblers to HMM-based approaches may in fact be an underestimate of their relative performance. Our discovery that predicted UTRs have high fractions of sequence that with BLASTX hits to NCBI proteins suggests that we failed to recover many CDS exons. Our pipeline for producing CDS annotations uses Transdecoder, and our results suggest that it may have a harder time correctly classifying CDS exons at the termini of a transcript than those within the transcript body. While long-read technology might help overcome this deficiency, we suggest that there is room for methods development to improve ORF detection from predicted CDS transcripts. Improved ORF detection and CDS exon boundary delineation would lead to improved performance with respect to several of the metrics we used to compare methods in this study.

Our findings also highlight the power and limitations of annotation transfer from another species with a high-quality annotation, as is done by TOGA. While TOGA often had high sensitivity (BUSCO scores) accompanied by low rates of intergenic predictions and gene fusions, we found that sensitivity could be much lower in plants, especially in those with larger genomes such as *Z. mays*. This undoubtedly stems from known difficulties performing whole-genome alignment with plant genomes (Song et al. 2024). Given the strong performance of RNA-seq assemblers across all the species we surveyed, researchers should consider RNA-seq assembly when there are known issues performing whole genome alignment for the species in question. For some taxa, there may be few if any closely related species with high-quality annotations, or genome alignments may be fragmented or contain many missing intervals in the target species.

Our finding that, for some species, there may complementarity among methods for which BUSCOs are recovered—and that RNA-seq based methods can recover genes that protein-based methods fail to predict—suggests that integration of annotations across multiple methods may potentially improve sensitivity. That MAKER performed poorly and TSEBRA appears to filter out real annotations when integrating protein and RNA-seq iterations of BRAKER, suggest that additional methods development on annotation integration is needed. Our naïve approach of consecutively adding annotations from one method that did not overlap with a base annotation also increased sensitivity, leading to an increase in the number of protein-coding genes with BLASTP hits to NCBI proteins. Nevertheless, we did not attempt to apply any filters to remove the spurious annotations that were invariably added in that workflow.

While we did not explore in depth how applying various filters impacted performance metrics, the frequent observation of high rates of intergenic predictions, as well as the occurrence of gene fusions, strongly suggest that more work is needed to identify filters that strike the balance between removing as many low-quality annotations as possible, while minimizing the filtering out of real sequences. It is often the case that filters applied in the literature appear *ad hoc*, even if guided by intuition and experience. We suggest a more quantitative approach is needed. For example, our application of random forest to classify genic and intergenic predictions suggests that machine learning approaches, while not perfect, offer great promise in identifying which variables should be used to set filtering thresholds. We found that whether a sequence had a BLAST hit to a set of known proteins, and expression level were useful discriminatory variables. Of course, a new genome does not benefit from the advantage of a “truth set” of real transcripts. However, a random forest (or other) model could, in principle, be trained with annotations from a related species, and that model could be applied to the predicted transcripts for a new genome assembly. Future work is needed to explore the utility of such an approach.

Even as much work remains to be done, our findings suggest some general guidelines for a researcher deciding how to annotate their newly assembled genome.

1. Generate RNA-seq data for at least the tissues related to the most pressing project needs, but ideally, across as many tissues as necessary to capture the species’ transcriptional complexity. Use Stringtie to assemble transcripts, and, until a better option is available, the Transdecoder workflow for adding CDS features to the annotation.
2. Consider using TOGA if RNA-seq data are not available and a well-annotated high-quality genome is available for a closely related species. If there is complementarity in the recovery of seemingly real protein coding sequences (determined with BUSCOs or gene symbols extracted from BLAST hits to established protein databases), consider an approach that integrates predictions of TOGA with BRAKER_protein_, giving more weight to TOGA that BRAKER.
3. If it is not possible to use TOGA and RNA-seq data are not available, use BRAKER_protein_.
4. If RNA-seq data are available and if there appears to be complementarity in the recovery of real protein-coding sequences between the methods, consider using an approach to integrate predictions from Stringtie and TOGA, giving more weight to Stringtie.
5. Either through a statistical approach such as random forest, or through heuristically-thresholded metrics (e.g. expression level, BLAST hits to an established protein database), remove predictions with a high likelihood of being intergenic.
6. Given the non-trivial frequency of fusions detected in the methods we analyzed (with the exception of TOGA), consider flagging genes that are likely fusions, e.g. if BLAST hits of different of different transcripts are to functionally distinct genes produced by genes with clearly different symbols, or similarly, if subsequences of individual transcripts have such divergent BLAST targets. Exclude these fusion genes from downstream expression analyses.
7. Consider using CGP in edge cases where, for example, there are a handful of incomplete annotations for some related species (perhaps generated by already available RNA-seq data), and the genome of interest is of small or modest size. Expect to do extensive filtering to exclude many spurious predictions.

In conclusion, the longer-term challenge for building genome annotations across the tree of life is to make methodological advances suggested above, and to integrate them into reproducible, automated workflows that can be deployed with minimal headaches for biologists. When this happens, population and comparative genomics studies will be easy to scale to hundreds, and even thousands of species, unleashing unprecedented power to tackle long-standing questions regarding the genetic architecture of phenotypic variation and the evolutionary mechanisms that generate and maintain biodiversity.

## METHODS

### Target taxa

Genome annotation tools are typically developed and optimized using high quality genome assemblies from a small suite of model organisms, e. g. *Homo* sapiens, *Caenorhabditis elegans*, and *Drosophila melanogaster*. As a result, it is difficult to generalize their performance in this narrow context to taxonomic groups that are highly divergent from those focal taxa, and for which the genome assemblies may not be of comparable quality. To facilitate more accurate generalizations regarding the performance of annotation methods, and, conversely, to explore whether there are effects of taxonomy and genome structure on annotation quality, we generated genome annotations for 21 species spanning six taxonomic groups: three species of heliconiine butterflies, three *Drosophila* species (dipterans), three birds, four mammals, four rosids, and four monocots (Supplemental Table 1). With the exception of the butterflies, we included as a “reference” a species for which both a high-quality genome assembly and annotation were available, and downloaded assemblies and annotations from NCBI. Each group contains at least one species that is relatively closely related to this reference. Because the NCBI genome versions and annotations for heliconiines are older than those widely used by the heliconiine research community, we used lepbase assemblies that were filtered to remove all scaffolds less than 1kb in length (Edelman et al. 2019). High quality annotations for these species were either unavailable, or generated by tools we evaluated and thus inappropriate to serve as a truth set. For example, the annotation for *H. melpomene* was generated with BRAKER. Because of the unavailability of gold standard annotations for heliconines, for these species we only generate a subset of performance metrics that don’t rely of an annotation truth set. Furthermore, *H. melpomene* is used as a reference assembly in whole-genome alignments (see below) but we did not generate annotations for this species.

### Genome assembly and reference annotation quality

To understand how features of genome assembly structure and quality might impact the quality of annotations, we generated several summary statistics describing genome contiguity and completeness. We generated standard statistics such as the total genome size, the contig and scaffold N50, and the fraction of genomic nucleotides that were soft-masked. We also quantified the number of single copy orthologs (BUSCOs) contained in a genome (Simão et al. 2015), and calculated a BUSCO score as 1 – (number missing BUSCOs/total number of BUSCOs searched).

To assess the quality of the reference genome annotations generated by NCBI, for each genome we extracted protein-coding transcripts and generated transcriptome BUSCO scores, and statistics on the minimum, maximum and median CDS length and fraction of CDS bases that were soft-masked. We also calculated the median RNA-seq alignment rate (see below) for each species’ annotation, as well as the fraction of transcripts for which estimated transcripts per million (TPM) was < 1.

### RNA-seq data acquisition and processing

For use with annotation methods that either build transcripts from RNA-seq reads or use read alignments to generate splice hints, for each species we downloaded 15-20 fastq file accessions from NCBI’s short read archive (SRA). We used the following criteria to choose accessions. We only considered 1) paired-end reads with a release data of 2011 or later, 2) at most one run per biosample, 3) Illumina reads sequence on HiSeq 2000, NextSeq or newer instruments (i.e. no Genome Analyzer II), 4) we excluded experimental treatments such as knockdowns, infections, and CRISPR modifications. If these criteria resulted in > 20 possible biosamples, we further required a minimum read length of 100bp, and for Metazoans, preferentially selected brain or head samples. If < 20 samples were available, we relaxed read length, release date, and instrument criteria with the goal of retaining 15 biosamples; with the exception of the heliconiine butterfly *D. plexippus* (for which 8 of 19 libraries had 36bp reads), we strictly excluded paired-end libraries where the read length was < 50bp. In a small number of cases where libraries contained hundreds of millions of reads, we down-sampled libraries to approximately 20 million read pairs with seqtk (https://github.com/lh3/seqtk).

To process the reads prior to sequence alignment, we stripped adapters with TrimGalore (https://www.bioinformatics.babraham.ac.uk/projects/trim_galore/). We did not trim low-quality bases at the ends of reads because the short read aligners used in this study soft-clip such bases such that they do not impact sequence alignment, and because such trimming can bias expression estimates (Williams et al. 2016). So as to avoid having to make inferences from SRA metadata regarding the strandedness of RNA-seq libraries, and to avoid the likely variable effectiveness of stranding protocols, we treated all libraries as unstranded, under the assumption that such information, if used would result in modest improvements in performance for annotation methods that leverage RNA-seq alignments.

### Annotation tools

#### RNA-seq assembly

We used RNA-seq reads to directly assembly transcripts with StringTie v. 2.1.2 and Scallop v. 0.10.5. These assemblers take as input spliced alignments of reads to the genome. We evaluated the impact of aligner by generating assemblies with two different aligners: HISAT2 v. 2.2.1 and STAR v. 2.7.9a. Following best practice, and to leverage evidence for splice sites across multiple samples, we used a 2-pass approach to generate STAR alignments. In the first pass, we generated an initial set of alignments for each sample. In the second pass, we concatenated the splice site tables generated for each sample’s first-pass alignment, and supplied the concatenated table as splice-site evidence for the second pass alignment of each sample. To generate a merged transcript assembly combining information across all individual-level assemblies, for StringTie and Scallop we use the stringtie-merge function and TACO (Niknafs et al. 2017) v. 0.7.3, respectively. With our heliconiine species, we initially evaluated two additional assemblers: PsiCLASS (Song et al. 2019) and Scallop2 (Zhang et al. 2022, 2). Poor performance relative to StringTie and Scallop, as well as excessive run times for PsiCLASS, led us to not consider these two tools further.

StringTie and Scallop annotations contain transcript and exon features: they do not predict CDS. Therefore, to incorporate CDS predictions into the merged annotations, we use a TransDecoder v. 5.5.0 (https://github.com/TransDecoder/TransDecoder) and an associated workflow (https://github.com/TransDecoder/TransDecoder/wiki) that predicts orfs, and then leverages orf predictions to predict CDS and UTR intervals associated with the gtf format input annotation file. After initial prediction of likely candidate orfs, we run blastp (Altschul et al. 1990) v. 2.12.0 searches against a protein database consisting of Uniprot and Trembl entries from all the species that we are attempting to annotate in the species group, e.g. for dipterans, the database consists of entries for *D. melanogaster*, *D. pseudoobscura*, and *D. yakuba*. We provide the search results as an input to transdecoder-predict, such that given two similarly scoring orfs, we preferentially keep the one with a blastp hit. In the interest of minimizing the filtering out of real orfs, we set the maximum e-value threshold for these blastp searches to 1 x 10^−4^. It should be noted that the workflow as described filters out of the final annotation any transcript without a retained orf prediction. The filtered orfs contain an unknown fraction of real orfs that TransDecoder failed to discover, as well as ncRNAs. While the focus of our research is on prediction of protein-coding genes, we provide as part of our code repository an accessory script for adding back into the final annotation these putative false negatives and ncRNA annotations.

#### Single-species ab initio methods

In contrast to transcript assembly-from-reads approaches, a long-established approach for predicting genes (and coding sequences in particular) is to parameterize hidden Markov models (HMMs) that are designed to traverse scaffolds, identify exon boundaries, and connect exons into transcript and gene-level features. The most sophisticated single-species versions of this approach use external evidence to parameterize HMMs and identify specific genomic locations where exon splice junctions are located. We evaluate BRAKER1 and BRAKER2 (both v. 2.1.6), which conduct iterative training and gene prediction using RNA-seq read and protein alignment evidence, respectively. Both BRAKER flavors wrap *ab initio* prediction with AUGUSTUS (Stanke et al. 2006) and GeneMark, with BRAKER1 using GeneMark-ET (Lomsadze et al. 2014) and BRAKER2 using GeneMark-EP+ (Brůna et al. 2020). Following developer recommendations, we provide protein evidence to BRAKER2 in the form of a protein fasta from OrthoDB v. 10 (Kriventseva et al. 2019) for the relevant taxonomic group, generated from pre-partitioned raw files as provided by the BRAKER developers (https://bioinf.uni-greifswald.de/bioinf), downloaded on 14 Septermber, 2018. For BRAKER1, we provide a bam file of RNA-seq STAR alignments merged across all libraries from the species being annotated.

While we consider multi-method annotation integration below, we also evaluate TSEBRA (Gabriel et al. 2021b), a python script-based tool to combine BRAKER1 and BRAKER2 runs, and select a well-supported subset of transcripts. We run TSEBRA following guidelines available at the TSEBRA github repository (https://github.com/Gaius-Augustus/TSEBRA), running it on the braker.gtf files (that include AUGUSTUS and GeneMark predictions) rather than on the AUGUSTUS-only annotations.

#### Single-species exon-aware liftover: TOGA

Using whole-genome alignments to transfer annotations across species from well-annotated to poorly or un-annotated species has a long history, e.g. with the UCSC Genome Browser LiftOver tool first becoming available in 2006 (Hinrichs et al. 2006). To perform such “liftovers”, we use TOGA (Kirilenko et al. 2023), which transfers CDS annotations across genomes in an exon-aware fashion that minimizes disruptions of ORFs. TOGA takes as input a whole genome alignment, and involves several steps, the details of which we provide at https://github.com/harvardinformatics/GenomeAnnotation-TOGA, and are an adaptation of the workflow described at https://github.com/hillerlab/TOGA. To remove potential spurious or bad annotation transfers, we filter out any transcripts in the primary annotation output (query_annotation.bed) for which there was not a corresponding entry in orthology_classification.tsv, i.e. transcripts for which TOGA could not determine an orthology class. Within each taxonomic group, we transfer annotations from the high-quality reference to all other species within the group, and from the species most closely related to the reference back to the reference species. For example, for dipterans we carry out three TOGA analyses, transferring *D. melanogaster* to both *D. pseudoobscura* and *D. yakuba*, and from *D. yakuba* to *D. melanogaster*.

#### Multi-species ab initio annotation

In studies seeking to perform phylogenetic comparative analyses, annotate multiple genome assemblies from related organisms, or where annotations or evidence (protein or RNA-seq) already exist for a subset of species of interest, methods that transfer evidence between lineages offer, in principle, a promising approach for performing genome annotation. We evaluate the most well-established approach for doing this, AUGUSTUS run in comparative mode (König et al. 2016), referred to hereafter as CGP. CGP relies on whole-genome alignment (WGA). Thus, as a first step, for each taxonomic group of genomes, we use Progressive Cactus (Armstrong et al. 2020) to produce a WGA. We then use an AUGUSTUS accessory script, *hal2maf_split.pl*, to split the hal-format cactus output file into multiple sub-files in multiple alignment (MAF) format; in doing so, we set as the “reference” genome (with which to provide coordinate anchors) a species with both a highly contiguous assembly and a high-quality annotation, and split in such a way so as to avoid splits that bisect the genomic coordinates of annotated genes in the reference. For each taxonomic group of species, we run CGP twice, once with splice site evidence from protein alignments, and once from RNA-seq alignments. For analysis with protein evidence, similar to analysis with BRAKER2, we use OrthoDB v.100 data representing the taxonomic group. For analysis with RNA-seq we used the merged STAR alignments across samples. In both instances, following guidelines from the developers (Hoff and Stanke 2019), we generate splice hints files for each species using scripts and code provided as part of the AUGSTUS package. In both modes, we do not predict UTRs, nor do we predict alternative isoform, i.e. one transcript prediction is made per putative gene. Detailed instructions regarding how we generated hints and run CGP are found at https://github.com/harvardinformatics/GenomeAnnotation-ComparativeAugustus.

#### MAKER

MAKER (Cantarel et al. 2008) is a genome annotation pipeline that has the ability to integrate multiple *ab initio* gene prediction packages, and to use protein and RNA-seq derived external evidence to perform post hoc curation of predictions. Because results with MAKER usually involve > 1 runs in order to retrain gene-prediction models, it is not a fully automated pipeline. Nevertheless, it has been used extensively due to its purported ease of use. MAKER also has the option to perform quality filtering and integration of annotations with EVidenceModeler (EVM) (Haas et al. 2008). For initial testing with three heliconiine species, we ran MAKER v. 3.01.03 four different ways that integrate predictions from AUGUSTUS (Stanke et al. 2006), SNAP (Korf 2004), and Genemark-ES (Lomsadze et al. 2005): 1) protein evidence only, without EVM; 2) protein and RNA-seq evidence, without EVM; 3) protein evidence only, with EVM; and 4) protein and RNA-seq evidence, with EVM. For protein evidence, we used the protein accessions associated with the lepbase (lepbase.org) Hmel2 genome assembly, which are proteins derived from BRAKER predictions. RNA-seq evidence was included as a gff3 file generated from the Stringtie assembly using STAR alignments of the species’ RNA-seq samples. We used default settings for the EVM configuration scoring file. To produce annotations, we ran MAKER twice closely following Daren Card’s detailed workflow (https://gist.github.com/darencard/bb1001ac1532dd4225b030cf0cd61ce2); see also https://github.com/harvardinformatics/GenomeAnnotation-Maker. Because with these test runs the use of EVM frequently produced lower quality annotations, and because we wished to evaluate the potential of MAKER as a full-service annotation tool, runs for other taxa were only performed with both protein and RNA-seq evidence and without EVM. Furthermore, because MAKER is computationally intensive and can take a considerable amount of time to run, for the other taxonomic groups we only generate MAKER annotations for the “reference” species of each group, with protein evidence being represented by the NCBI protein accession associated with the genome for the next closely related species in the set of species we annotated within each taxonomic group.

### Annotation quality metrics

Accurate annotation of UTRs is challenging, and even more so for non-model organisms for which RNA-seq data are typically sparse. Similarly, long non-coding RNAs are also difficult to annotate (Uszczynska-Ratajczak et al. 2018). In most contexts, neither feature is crucial to genome-enabled evolutionary studies in non-model organisms. Thus, we focus our evaluation of genome annotation methods on protein coding sequences, filtering out UTRs and non-coding loci. For annotation methods that include the UTR portions of mRNAs, we strip UTR exons (and any UTR-labelled features) from annotation gtf or gff3 files. We also do this for NCBI gff3 files prior to comparisons with the annotation methods we evaluate.

We use the NCBI genome annotations as putative sets of “true” annotations. Although the quality of these annotations certainly vary due to many factors – the quality of the genome assembly, the amount and kind of experimental evidence available at the time the annotation was generated, challenges in annotating larger, more complex genomes—we believe they are a reasonable approximation to annotations one would hope to achieve with a new genome assembly, using stand-alone annotation tools deployed on local HPC clusters. Nevertheless, while we make comparisons to NCBI annotations for all species-tool combinations, we pay particularly close attention to those species for which we know the genomes and annotations are of the highest quality: *H. sapiens*, *D. melanogaster*, *Z. mays*, *Arabidopsis thaliana*, and *Gallus gallus*.

Details on bioinformatics package command lines and custom python scripts are available in the relevant GitHub repositories detailed in the DATA ACCESS section at the end of this paper.

#### Annotation completeness: transcriptome BUSCOs and expression

To assess transcriptome completeness, we calculate transcriptome BUSCO scores and compare them to scores for the NCBI transcriptomes. For comparisons across all species-method combinations, and in order to normalize for varying degrees of genome assembly completeness and quality, we calculate ratios of transcriptome BUSCO score to that for the respective genome. To evaluate the extent to which the predicted transcriptome represents the expressed transcriptome, we then use RSEM (Li and Dewey 2011) v. 1.3.3 to wrap bowtie2 (Langmead and Salzberg 2012) alignment of each RNA-seq library to the predicted transcriptome, and estimate both gene and isoform-level expression. From those alignments, for each annotation we calculate the median alignment rate (across the set of samples), and the proportion of genes and transcripts for which TPM < 1. Because using the same RNA-seq libraries to generate transcriptome assemblies with Stringtie and Scallop may bias alignment rates upwards relative to tools that don’t leverage evidence from those RNA-seq libraries, we also perform alignments on an additional test set of six RNA-seq paired-end SRA accessions for each of our five reference species.

#### Protein-level statistics

For each genome annotation we report the number of protein-coding genes, and the number of CDS transcripts. We use GetProteinFastaStats.py, a custom python script that leverages biopython (Cock et al. 2009) modules to calculate mean and median protein lengths, the fraction of predicted proteins that have internal stop codons, and the fraction that are complete, where complete is defined as having proper start and stop codons, and no internal stop codons. For TOGA annotations, we generate these statistics from the protein sequences the software outputs (after filtering out those without ortholog classifications). For all other annotation tools we used the version of gffread distributed with cufflinks (Trapnell et al. 2010) v. 2.2.1, to extract the protein sequences.

To estimate the fraction of protein predictions that were real, we used blastp (Camacho et al. 2009) v. 2.12.0 to search for matches, with a maximum e-value of 1 x 10^−5^, against a database consisting of the NCBI protein accessions for all of the species that were used in our study for the taxonomic group of interest.

#### False positives: intergenic predictions

Motivated, in part, by some tools predicting far more CDS transcripts that are recorded in NCBI annotations, we assessed whether this could be due to (presumably incorrect) intergenic predictions, where intergenic is defined as falling entirely outside of the CDS intervals for all protein-coding genes annotated by NCBI. To do this, for each new annotation, we generate transcript interval and gene interval bed files, where each entry represented the genomic boundaries of the transcript or gene (excluding UTRs), respectively. We then use bedtools v. 2.26.0 (Quinlan and Hall 2010) to intersect these files with a bed file consisting of UTR-stripped NCBI gene boundaries, recording the number of bases of overlap such that only same-strand overlaps were counted as overlaps, e.g. *intersectBed -s -wao -a newannotation_intervals.bed -b NCBI_gene_intervals.bed*. We then count the number of predicted transcripts and genes lacking any overlap with NCBI gene coordinates.

#### Gene fusions

Real fusion events in which a transcript contains CDS from different annotated genes should be extremely rare. Thus, the presence of non-trivial frequencies of predicted transcripts for which exons span multiple NCBI genes most likely represent bioinformatics pipeline-induced errors. We evaluate fusions at the gene level for which we define three types: 1) an individual predicted CDS overlapping with CDS from multiple NCBI genes, 2) different CDS originating from the same predicted gene overlapping with the CDS of different NCBI genes, and 3) cases where both of these types of fusions occur. To quantify the frequency of these fusion events, we first converted the CDS features of a species’ NCBI annotation and the CDS annotations of a method being evaluated to bed format. Next, we used bedtools to perform a “left outer join” of NCBI CDS features to those of the new annotation, e.g. *bedtools -loj -a newannotation_cds.bed -b ncbi_cds.bed > loj_overlaps.bed.* We then evaluate each gene in the new annotation relative to the first two classes of fusions using a custom python script, BuildProteinCodingGeneCdsFusionSummaryTable.py and including the -filter-nested switch that excludes from calculations known fusions that are present in the NCBI annotation. We calculate the frequency of all three classes of fusions using basic awk commaxnds. Our bedtools operation does not enforce same-strand matching, but our python script does, such that we do not consider overlaps between new predictions and NCBI predictions on opposite strands.

#### Undetected CDS in UTRs

Our RNA-seq assembly pipelines integrate ORF finding with Transdecoder such that exon features are decomposed further into CDS and UTR features. While we exclude UTRs to make assembler performance metrics comparable to tools that do not predict UTRs, we observed a substantial drop-off in RNA-seq read alignment rates to the Stringtie and scallop annotations when UTRs were filtered out: unfiltered annotations had much higher alignment rates than all other methods, but were on par with those methods after excluding UTR intervals. To examine the cause of this phenomenon, we considered three possible causes. First, because for relatively complete high-quality annotations the NCBI annotation pipeline will computationally truncate UTRs to prevent stop-codon readthrough, we contrasted the length distributions RNA-seq assembler UTRs and those from NCBI, expecting that the disparity would be greater for the reference genomes for each of our taxonomic groups than for other, more recent genome assemblies for which truncation would not be as severe. Under this scenario, the reduction in alignment rate after UTR removal would be due to an excess of reads originating from transcriptional readthrough (or because the NCBI UTR truncation was overly conservative). Next, we considered the possibility that Transdecoder consistently fails to predict CDS orfs at the terminal ends of transcripts, such that a large proportion of real CDS sequences is being incorrectly filtered out when we strip out UTRs. To test this hypothesis, we extracted the UTR sequences from the Stringtie and scallop annotations and used blastx (Camacho et al. 2009) v. 2.12.0 to search for matches against the NCBI protein sequences from the same species’ NCBI accession, with a maximum e-value of 1 x 10^−5^. We calculated the fraction of transcripts for which at least one of the UTRs had a hit to the protein database.

#### Mulit-method integration

To determine whether integration of multiple annotation methods could harness the complementary strengths of individual methods, we tested an approach where we 1) set one annotation as the “base annotation”, 2) add features from a second annotation methods that fall entirely outside of the gene intervals for the base annotation, and 3) iterate this process for additional methods, e.g. adding features from a third method that do not overlap genes from the integration of the first two annotations. We demonstrated this approach for *Homo sapiens*, investigating the effects of different choices for base annotation, exclusion of RNA-seq assembler or TOGA, and whether there are decreasing returns with increasing number of integrated methods. For these comparisons we focused on four metrics: BUSCO score, RNA-seq alignment rate, the total number of genes with BLASTP hits to NCBI reference proteins, and the percentage of all integrated genes that have such hits. While we do not compare this approach directly to the other integration methods evaluated here (MAKER and TSEBRA,) our results demonstrate that both of these annotation tools under-perform relative to other standalone methods.

## DATA ACCESS

Detailed explanation of steps in annotation pipelines, along with associated python scripts for data processing are provided in several repositories listed below

RNA-seq transcript assembly: https://github.com/harvardinformatics/GenomeAnnotation-RNAseqAssembly/
BRAKER: https://github.com/harvardinformatics/GenomeAnnotation-Braker
TOGA: https://github.com/harvardinformatics/GenomeAnnotation-TOGA
CGP: https://github.com/harvardinformatics/GenomeAnnotation-ComparativeAugustus
MAKER: https://github.com/harvardinformatics/GenomeAnnotation-MAKER

In addition, HPC slurm job scripts and specific command lines used to run annotation tools in this paper, as well as python scripts to generate annotation quality metrics are available at https://github.com/harvardinformatics/GenomeAnnotation.

## Supplementary Materials

**Figure S1.**
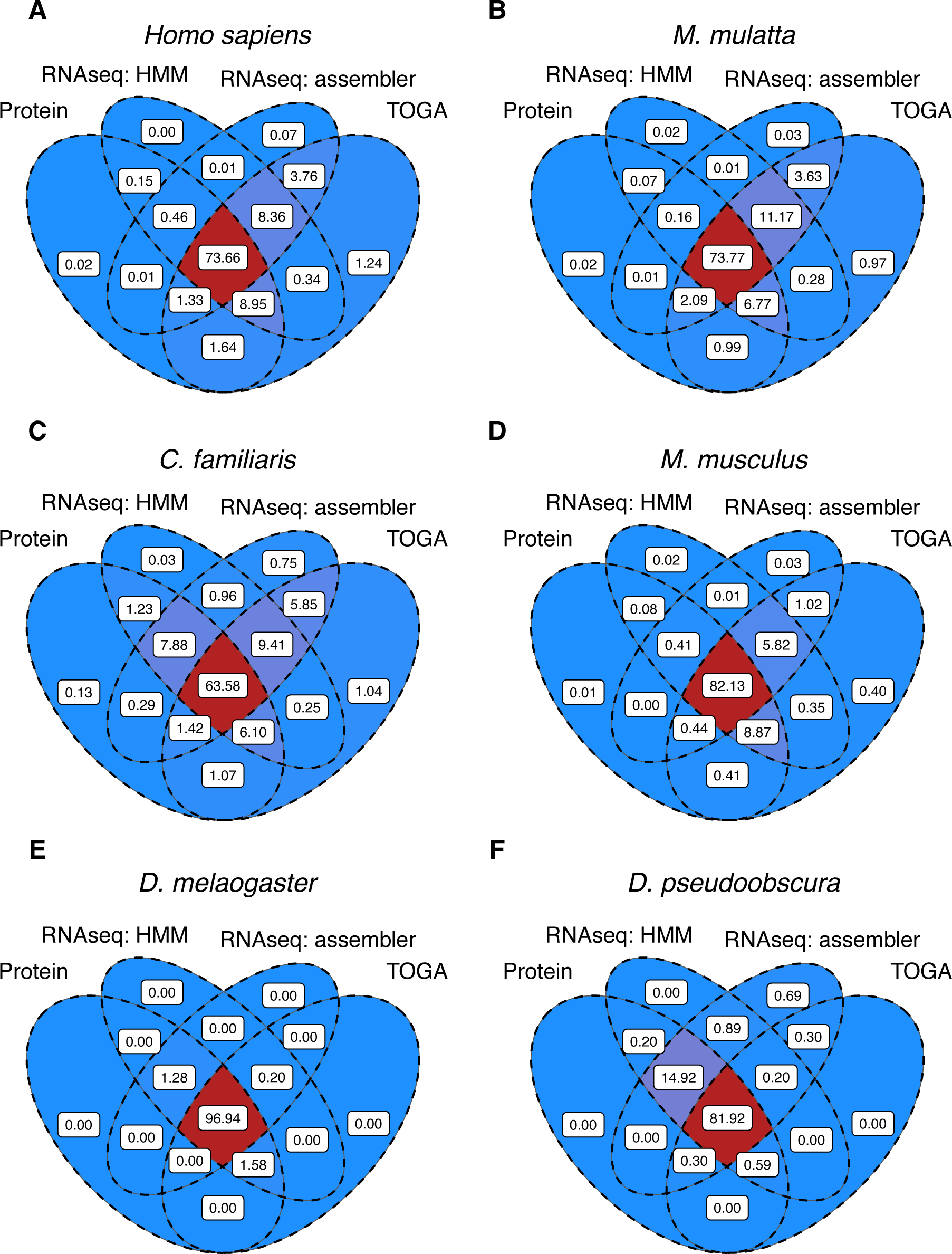
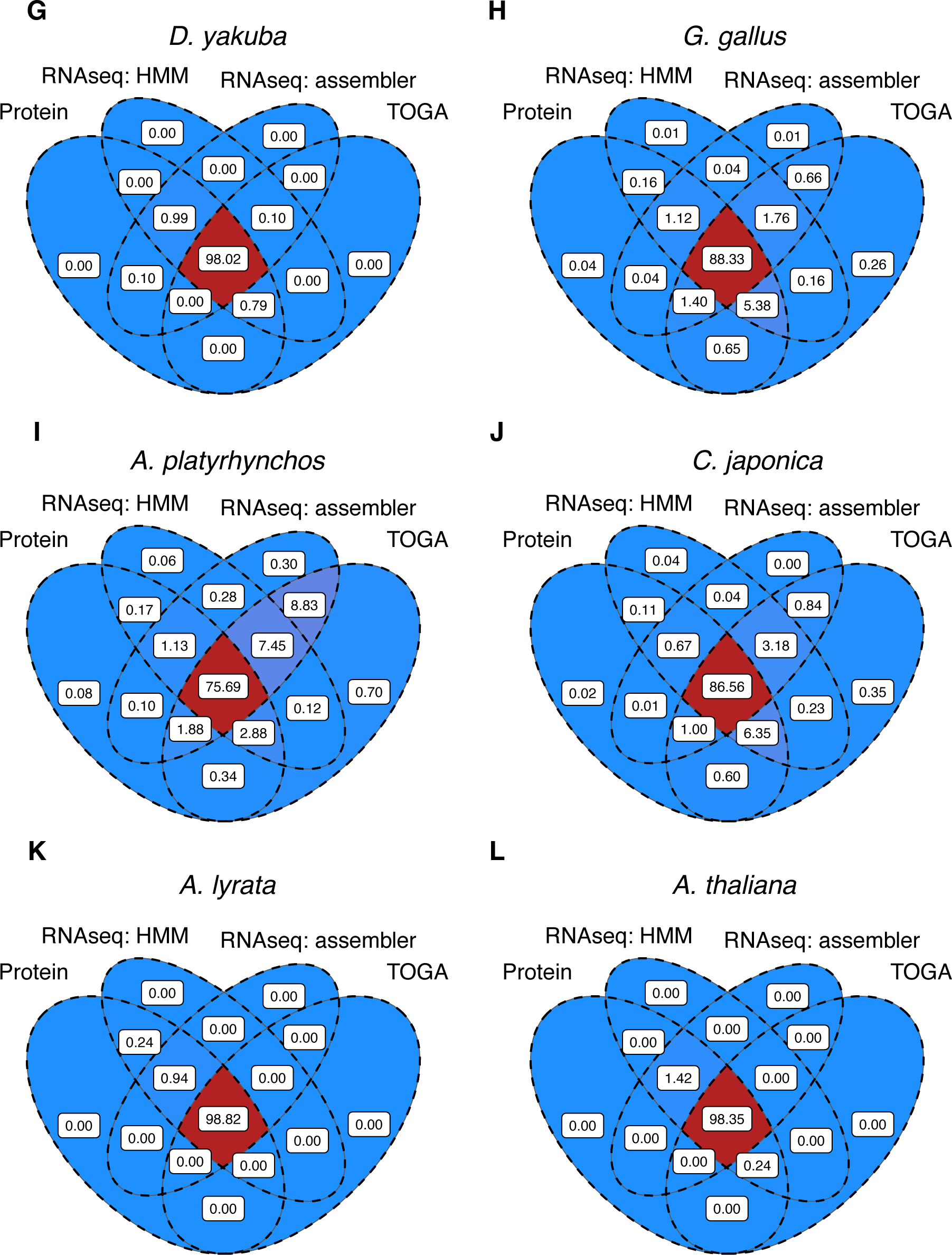
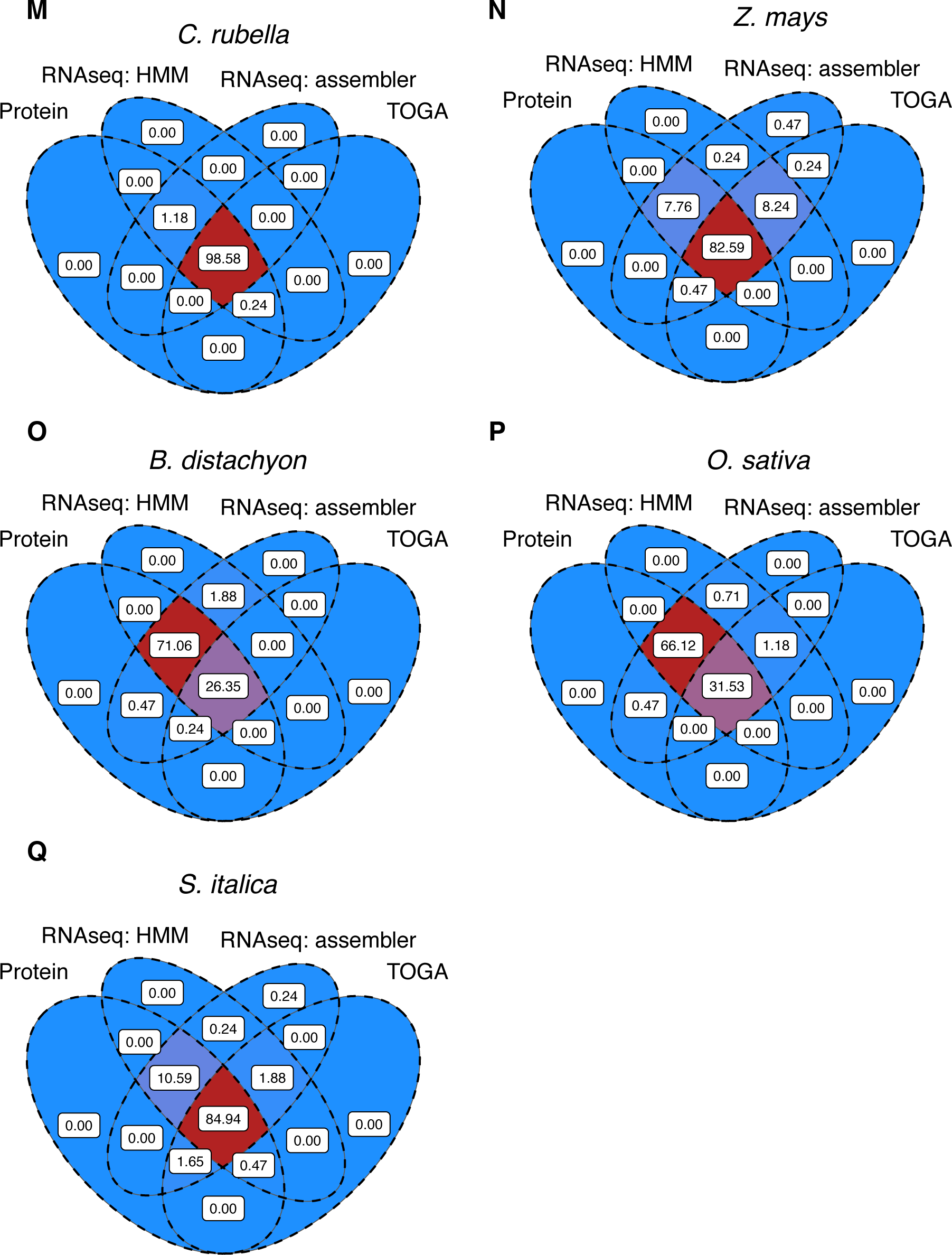
Percent of recovered BUSCOs that overlap between methods for species in addition to Figure 2. (A-D) mammals, (E-G) dipterans, (H-J) birds, (K-M) rosids, and (N-Q) monocots.

**Figure S2.**
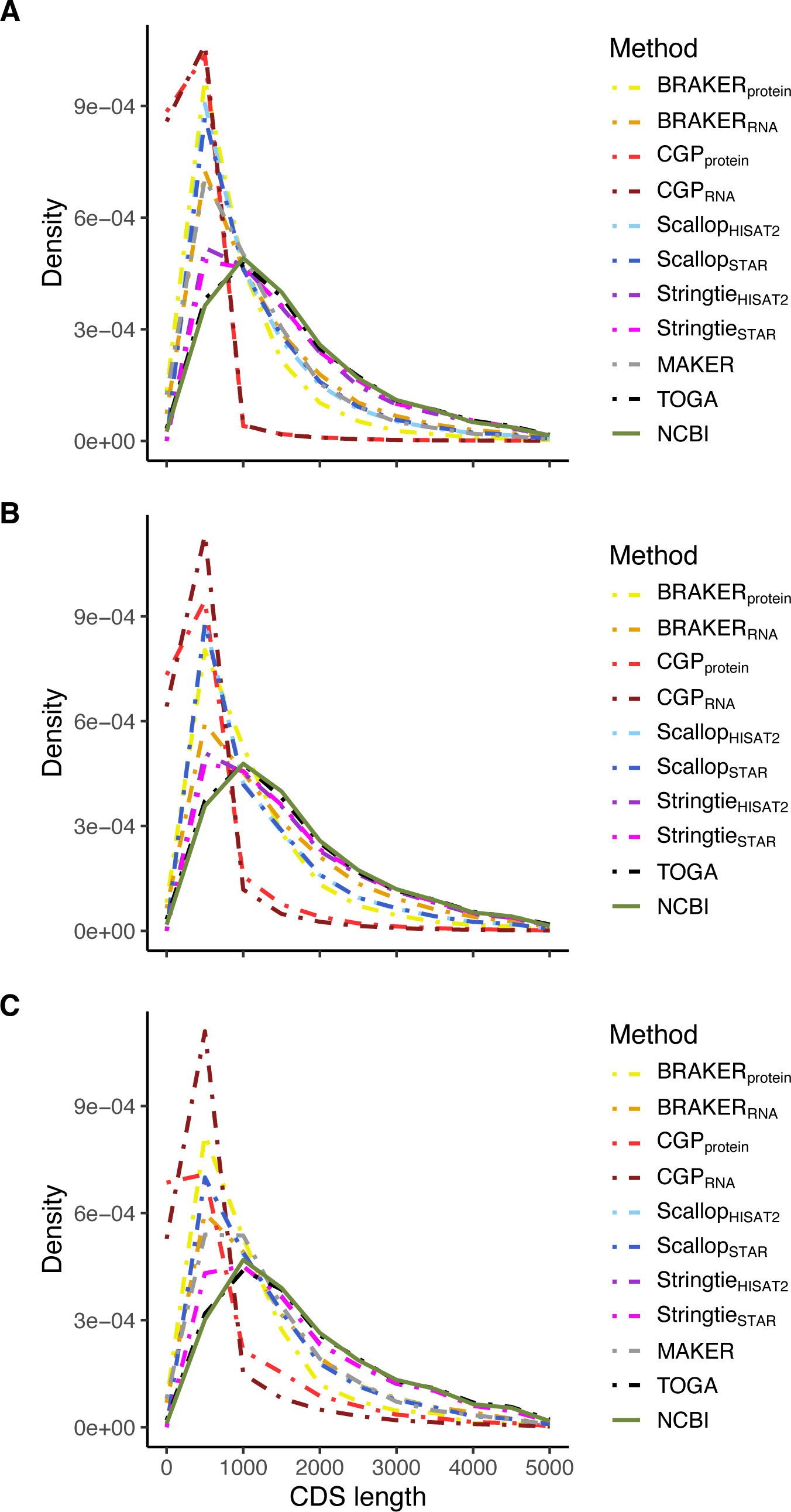
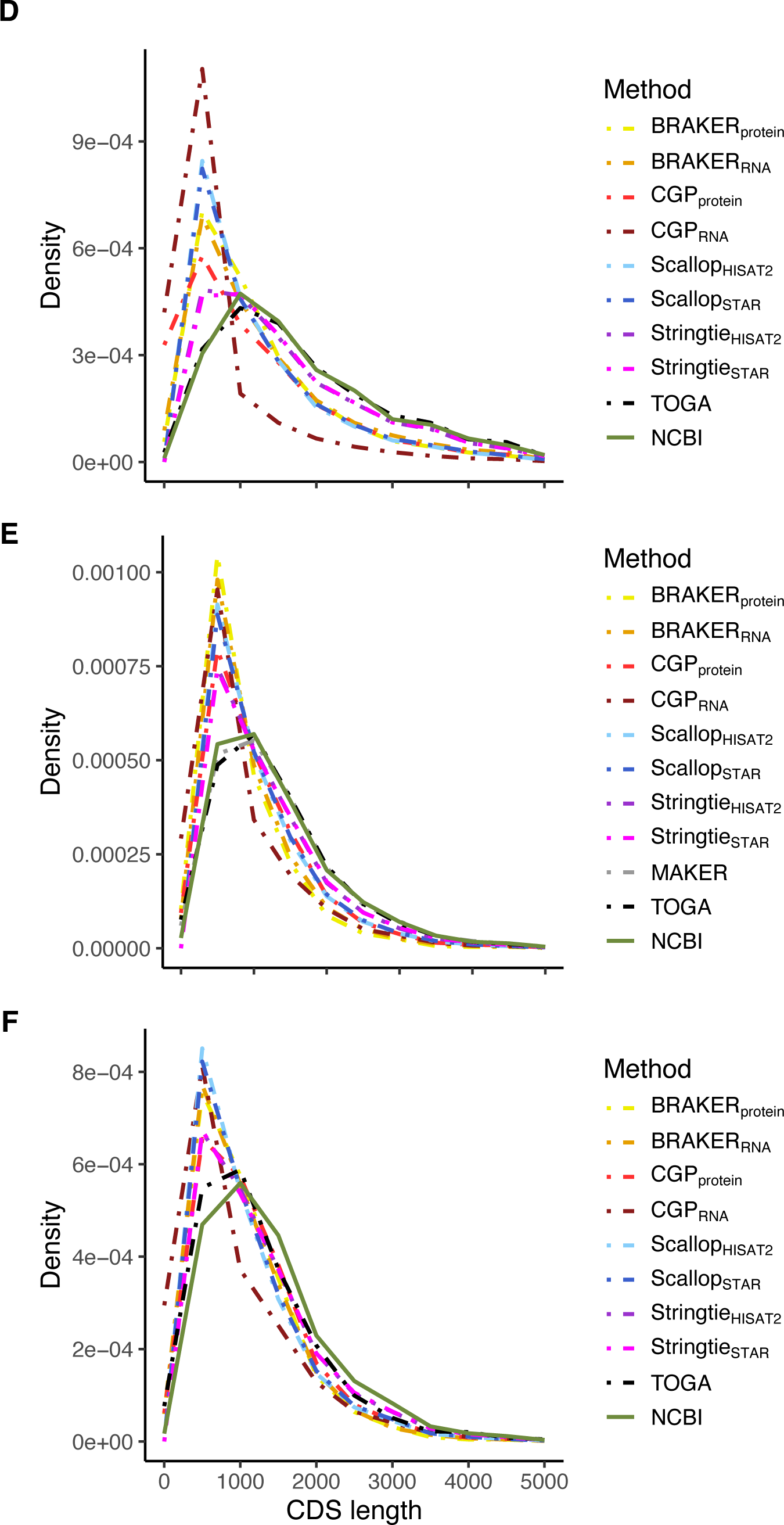
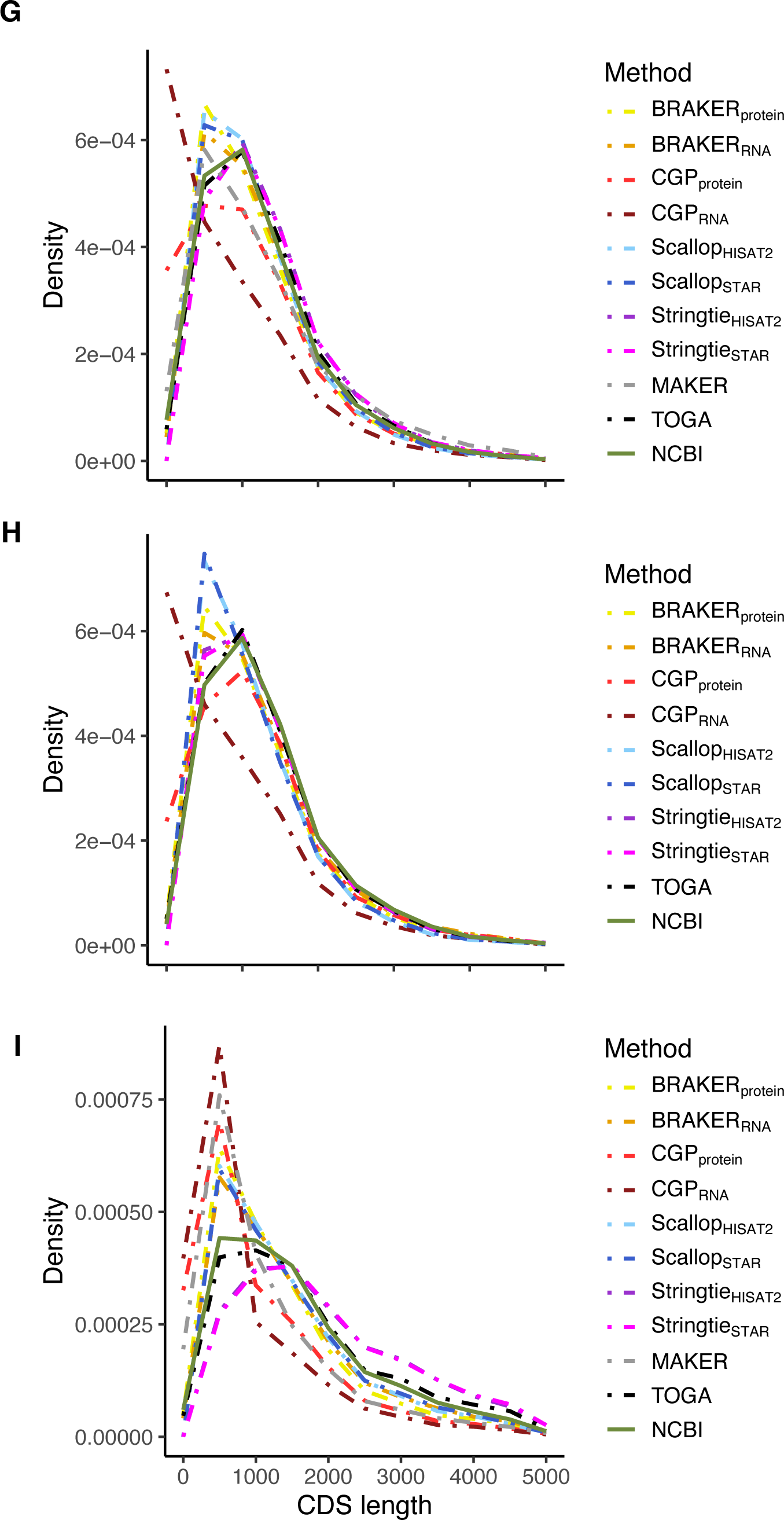
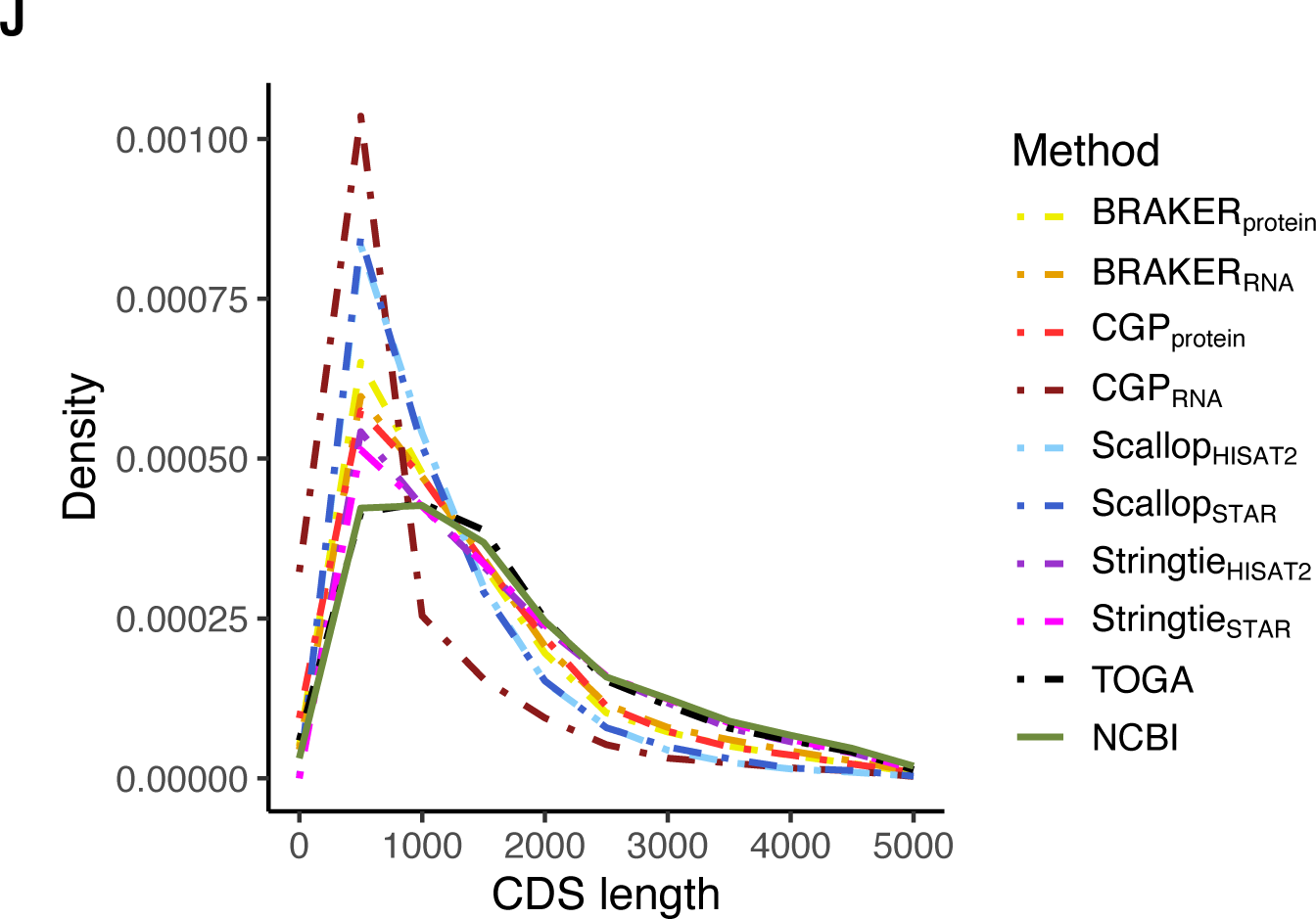
CDS length distributions for annotation methods and NCBI benchmark for (A) *H.* sapiens, (B) C*. familiaris*, (C) *G.* gallus, (D) *A. platyrhynchos*, (E) *Z. mays* (F) *O. sativa*, (G) *A. thaliana*, (H) *C. rubella*, (I) *D. melanogaster*, and (J) *D. pseudoobscura*.

**Figure S3.**
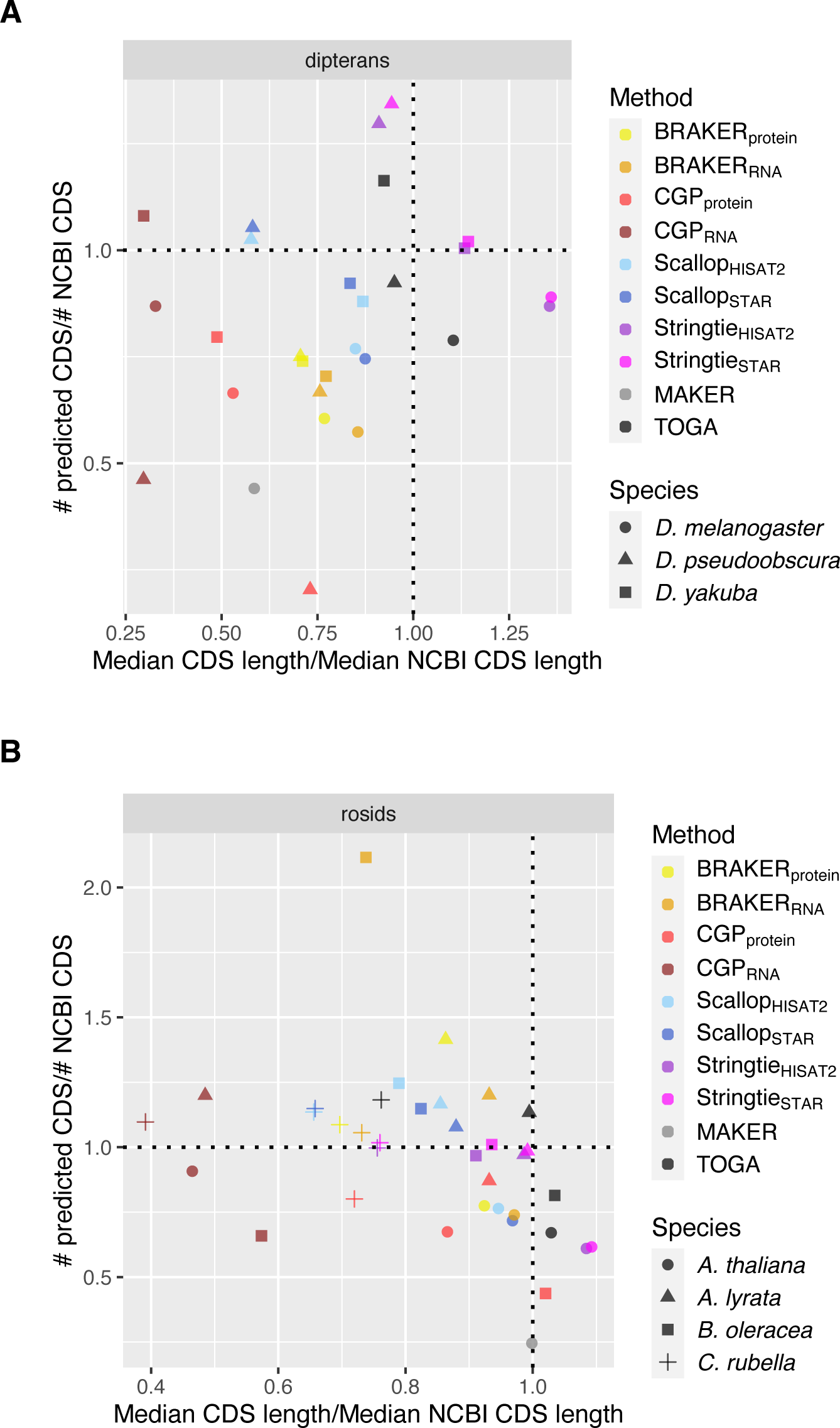
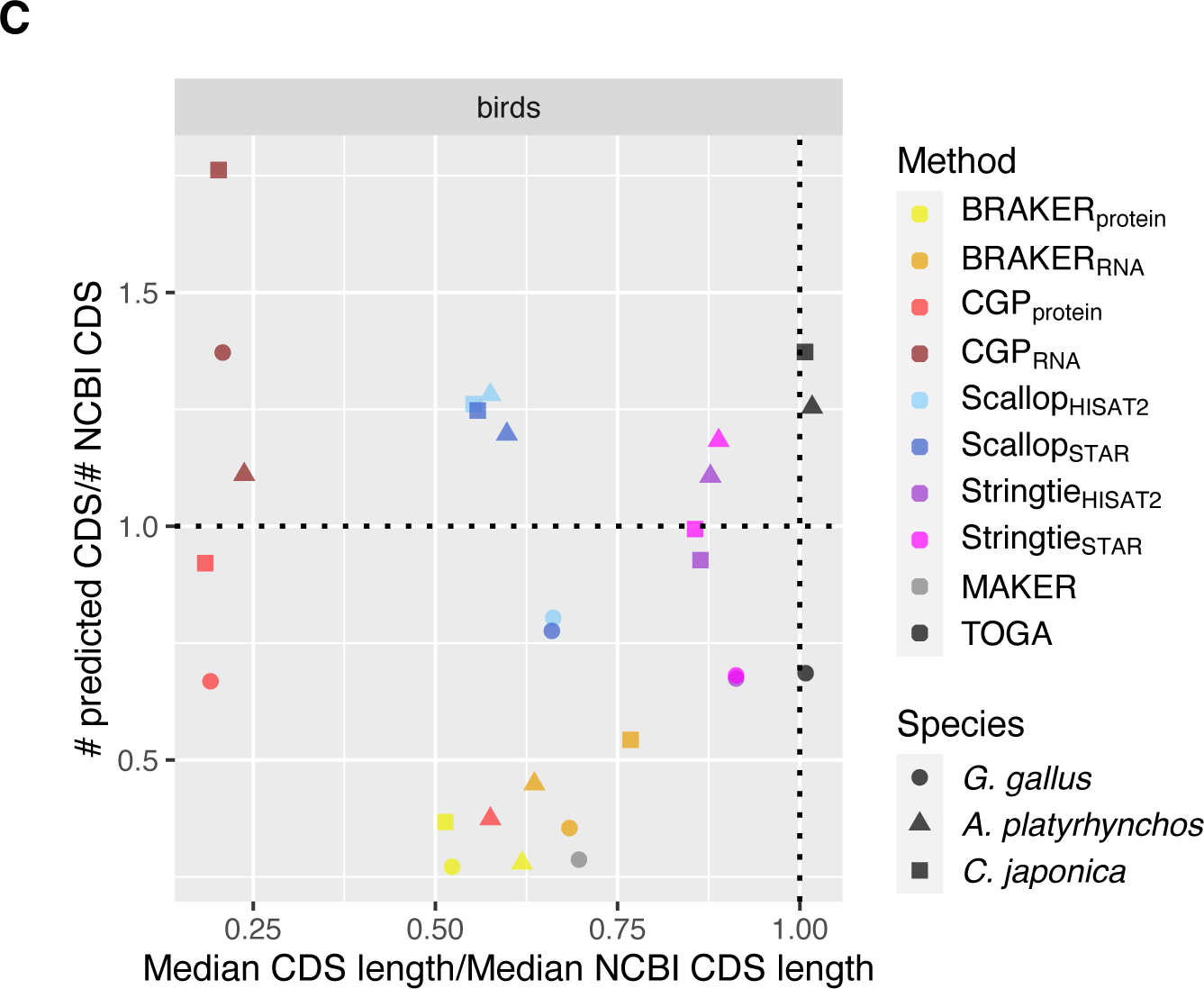
Joint distributions of number of predicted CDS (normalized by number of NCBI predictions) over median predicted CDS length (normalized by median NCBI CDS length) for (A) dipterans, (B) rosids, and (C) birds. Dotted lines indicated equivalence to NCBI annotation, such that methods that are closest to the intersection of those lines best approximate CDS length and number of NCBI annotations.

**Figure S4.**
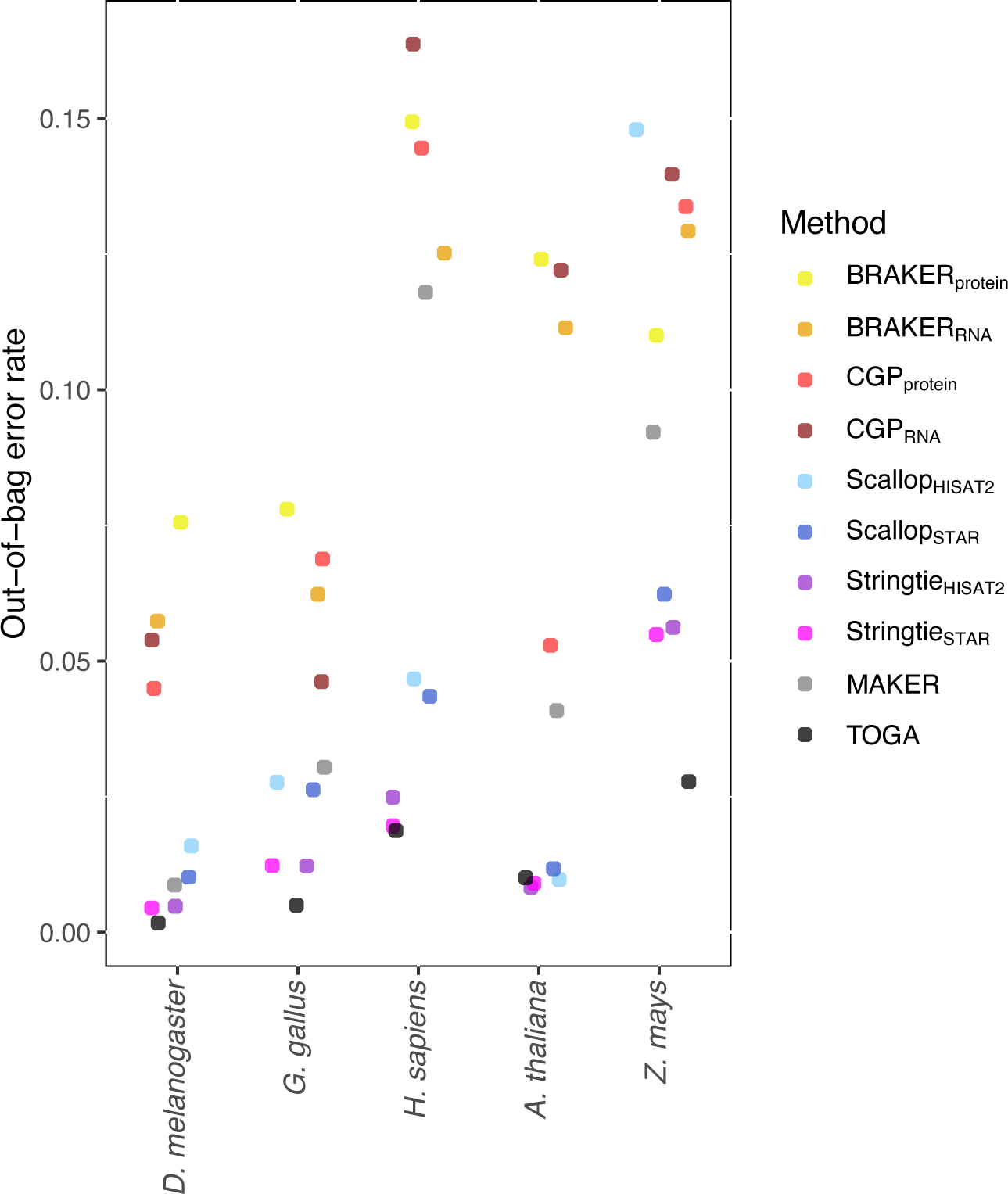
Out-of-bag error rates for random forest models classifying CDS transcripts as either overlapping NCBI gene intervals, or falling outside of those intervals, i.e. intergenic. Models are generated for the reference species for each taxonomic group for which annotations are thought to be complete or nearly so.

**Figure S5.**
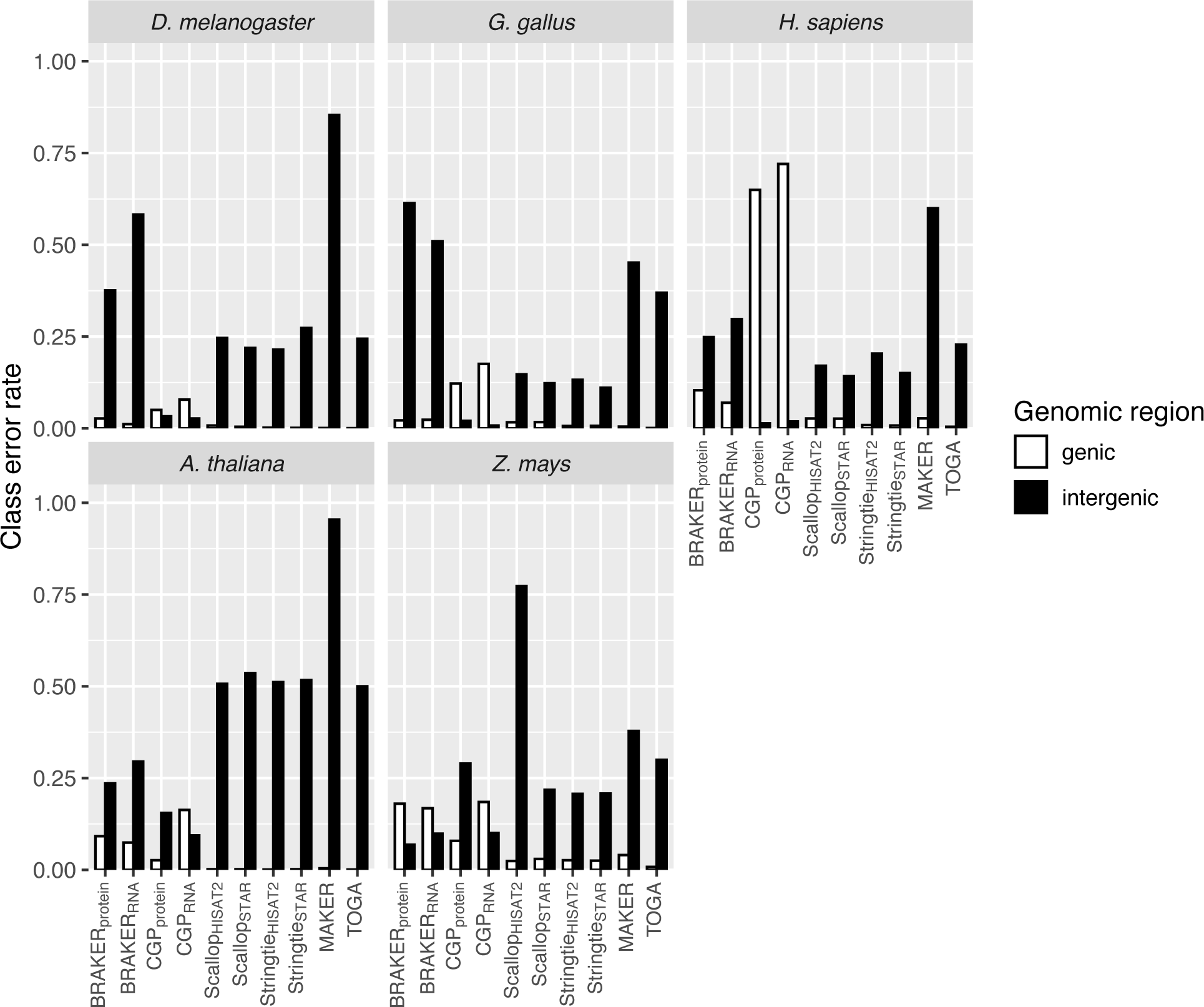
For reference species, out-of-bag error rates broken down by predicted class, for random forest predictions of CDS overlapping NCBI protein-coding genes, vs. those entirely outside of NCBI protein coding gene intervals.

**Figure S6.**
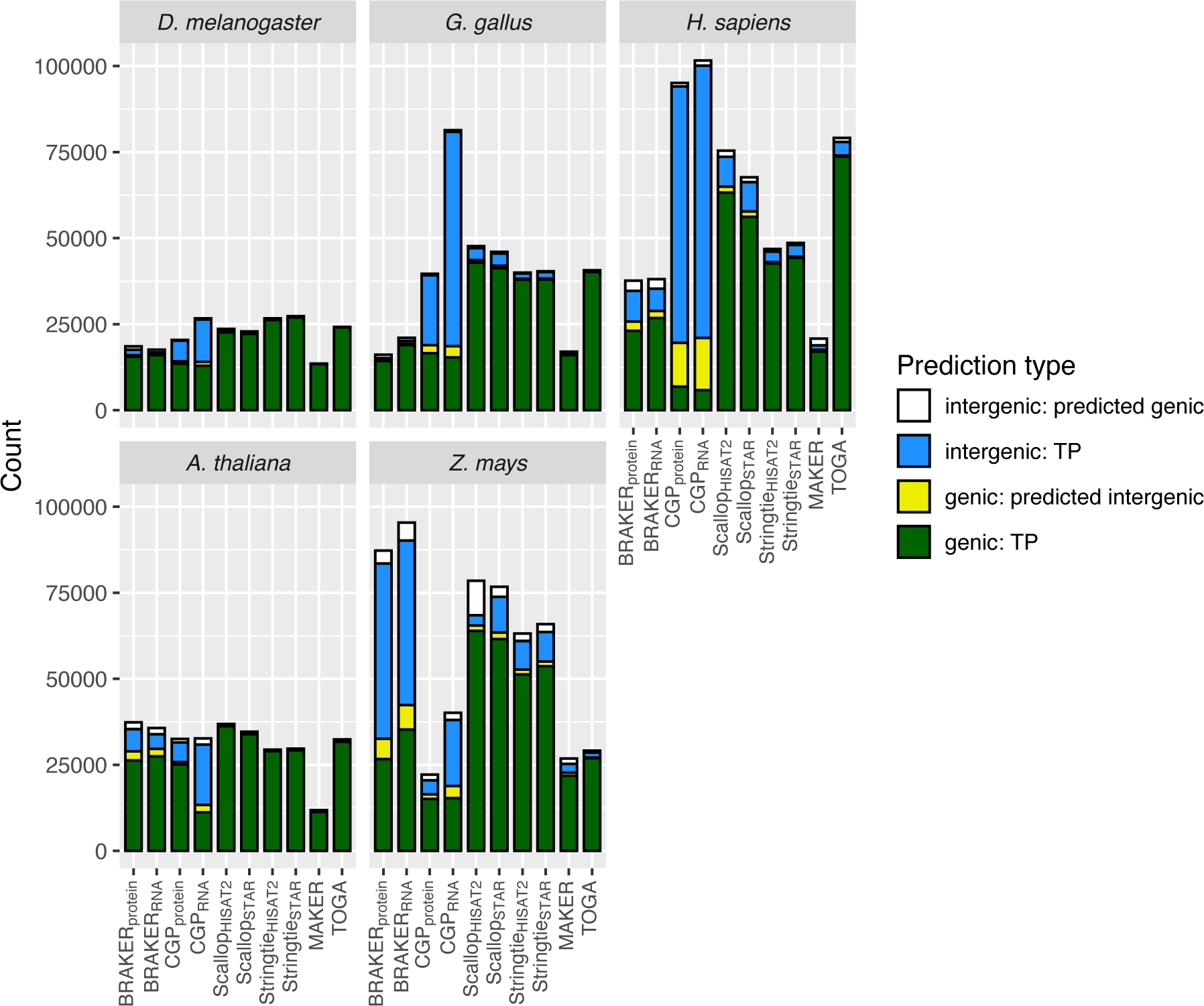
Random forest confusion matrices for reference species, for prediction accuracy of models predicting the genic versus intergenic status of CDS transcripts based upon transcript features.

**Figure S7.**
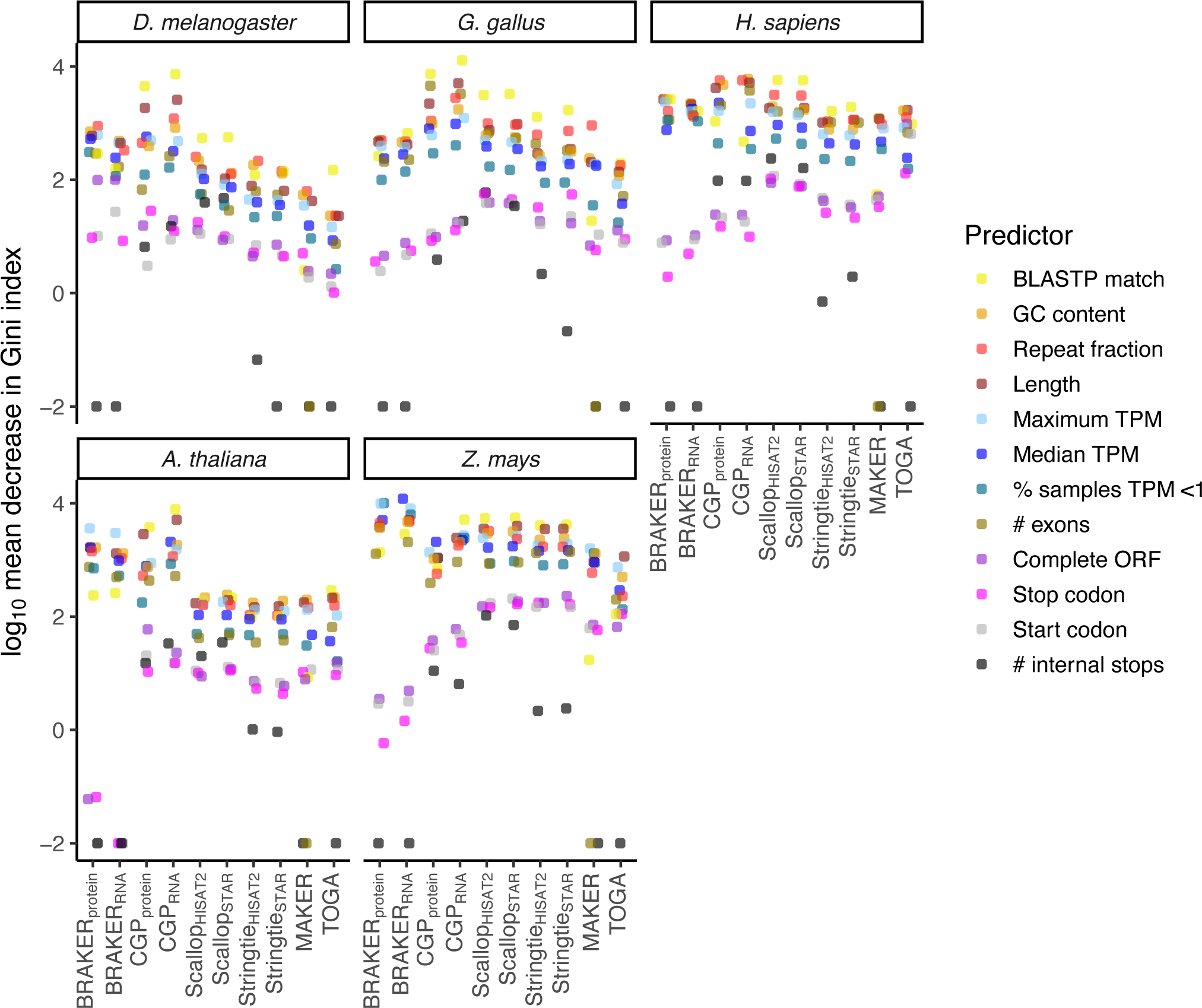
Decrease in Gini index, an estimator of variable importance for random forest predictions of intergenic versus genic region location of predicted transcripts for reference species.

**Figure S8.**
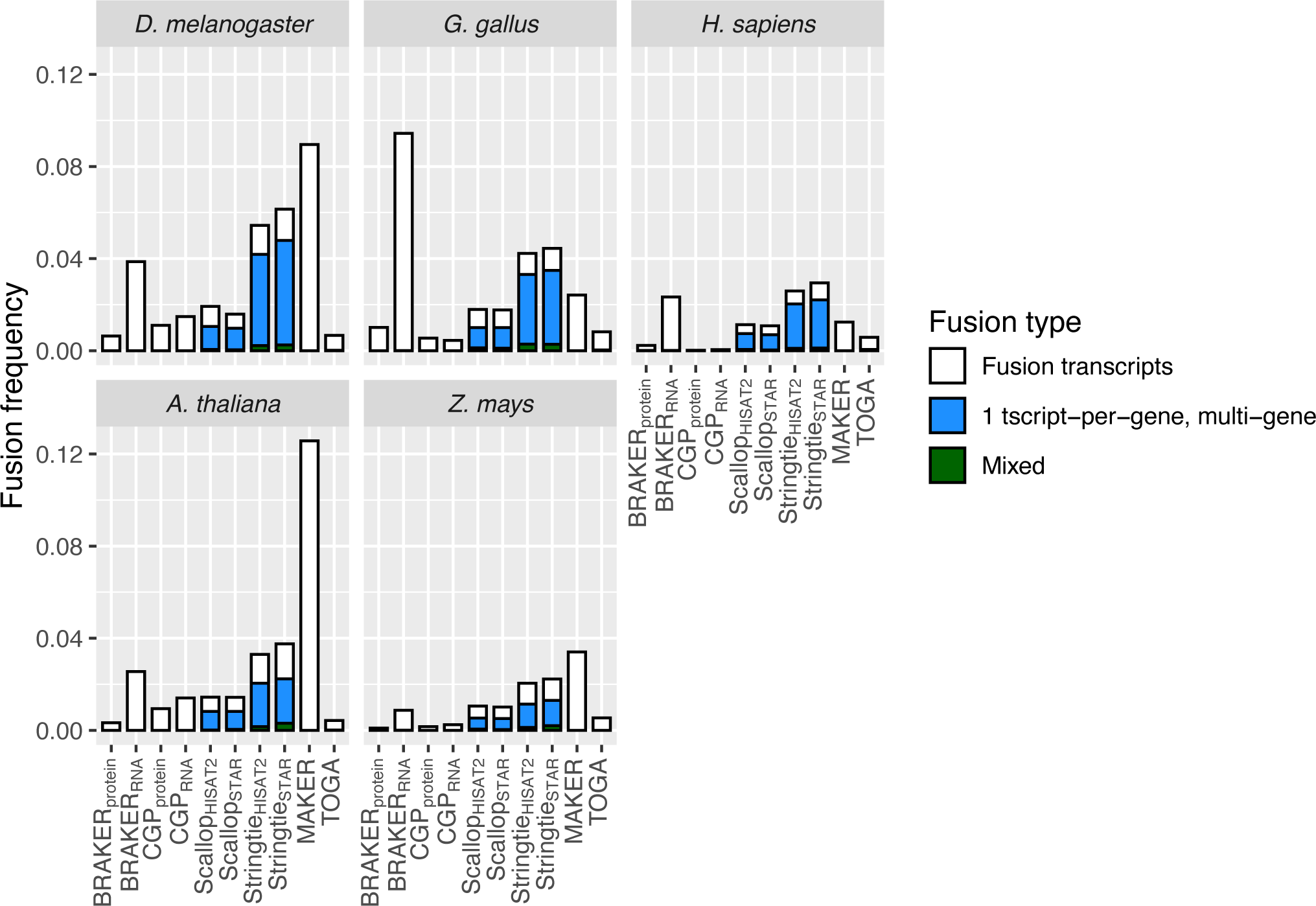
Frequency of gene fusions, broken down by fusion type for five reference species Fusions are defined as when individual transcripts overlap the CDS of multiple NCBI genes (fusion transcripts), different transcripts from the same predicted gene each have overlaps to different genes (1 tscript-per-gene, multi-gene), or a combination of both of these (mixed). Frequencies are calculated after filtering out NCBI-annotated fusion events.

**Figure S9.**
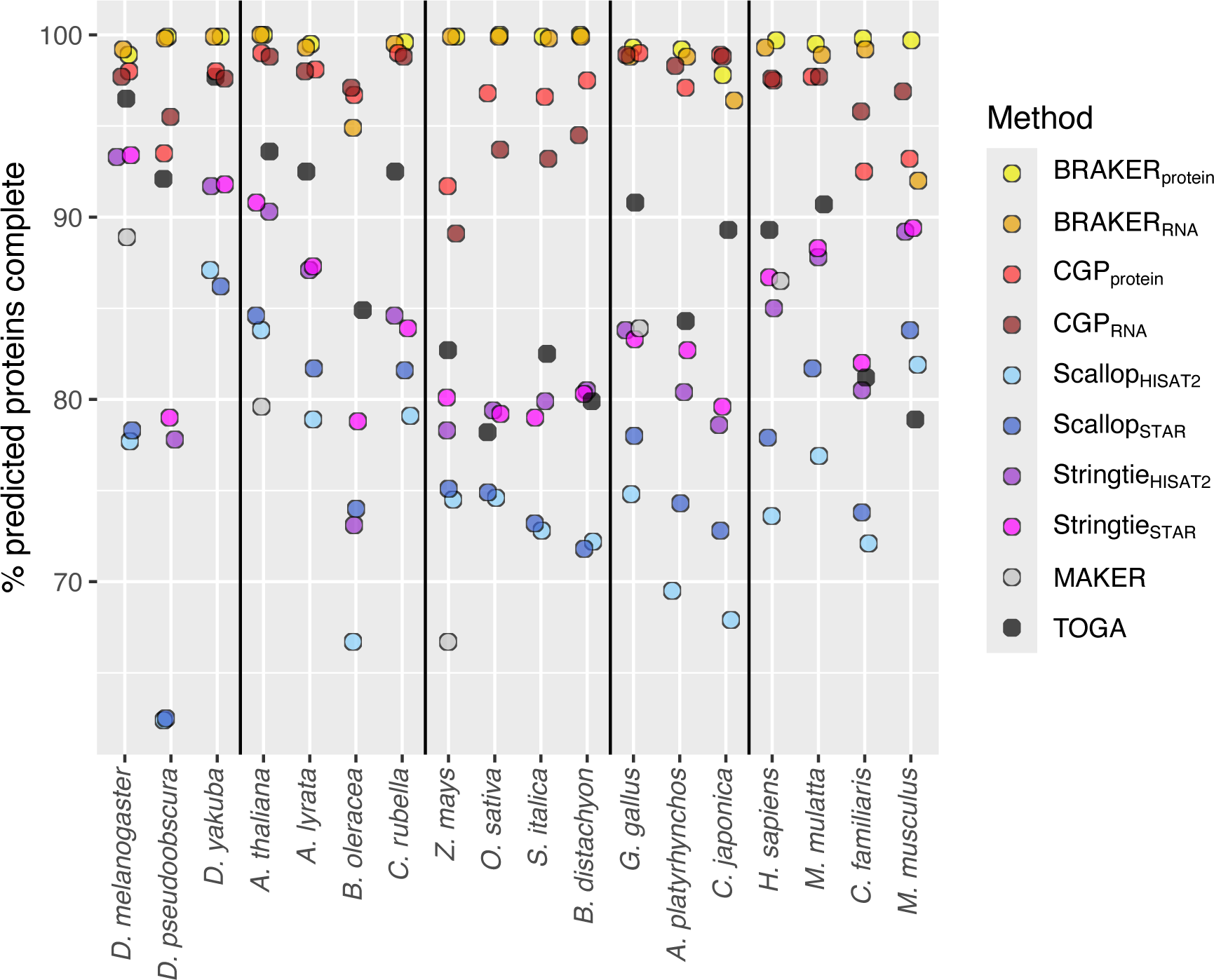
Grouped by species and method, percentage of predicted proteins that are complete, having both initiating start and terminating stop codons, and no internal stop codons.

**Figure S10.**
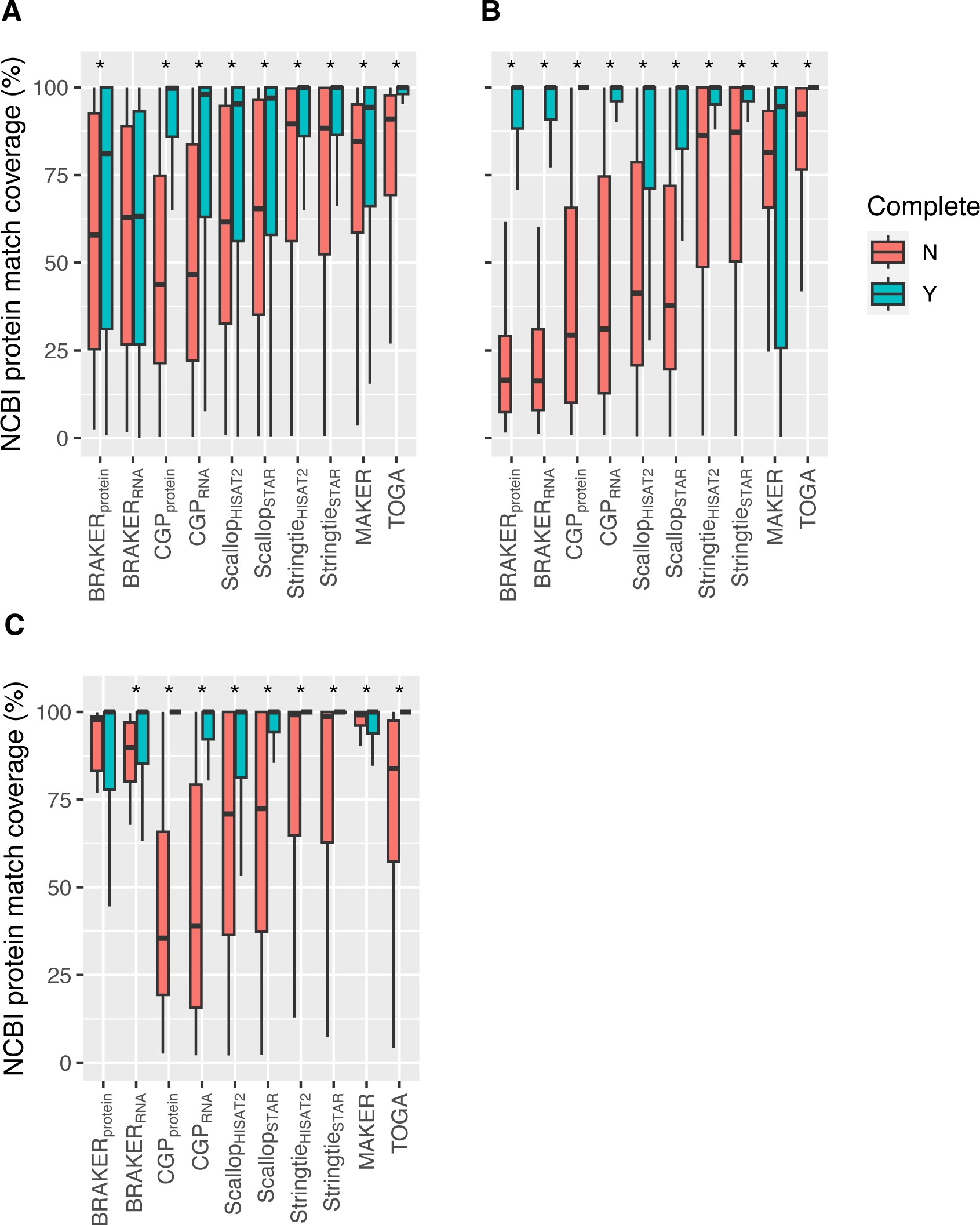
For predicted proteins with BLASTP hits, proportion of NCBI best hit target covered by amino acid matches with the predicted protein for (A) *G. gallus*, (B) *D. melanogaster*, and (C) *A. thaliana,* broken into proteins that are complete (start and stop codons present, no internal stop codons), and those that are not. Benjamini-Hochberg adjusted p-values for Wilcoxon rank-sum tests, p≤0.05 indicated with *.

**Figure S11.**
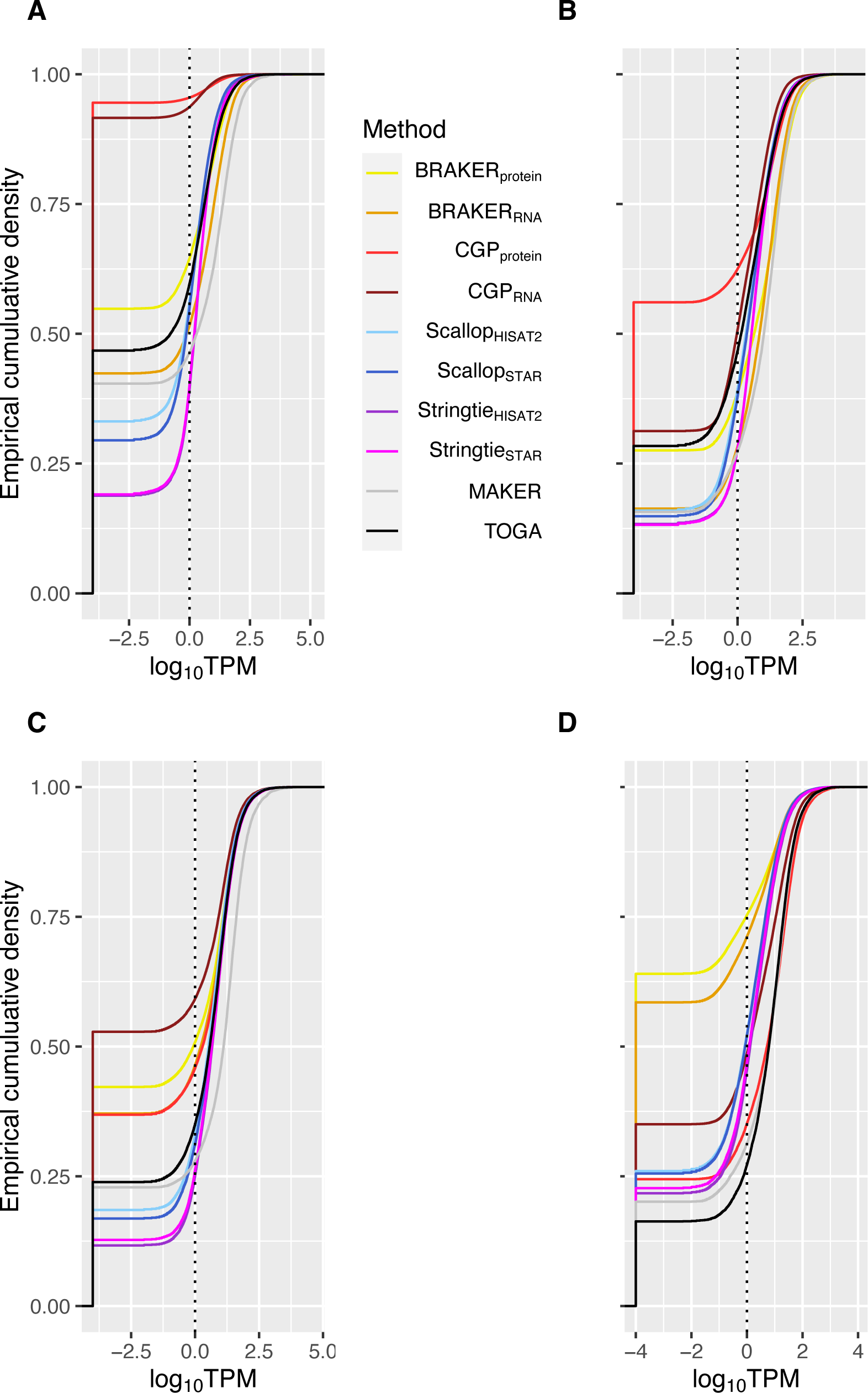
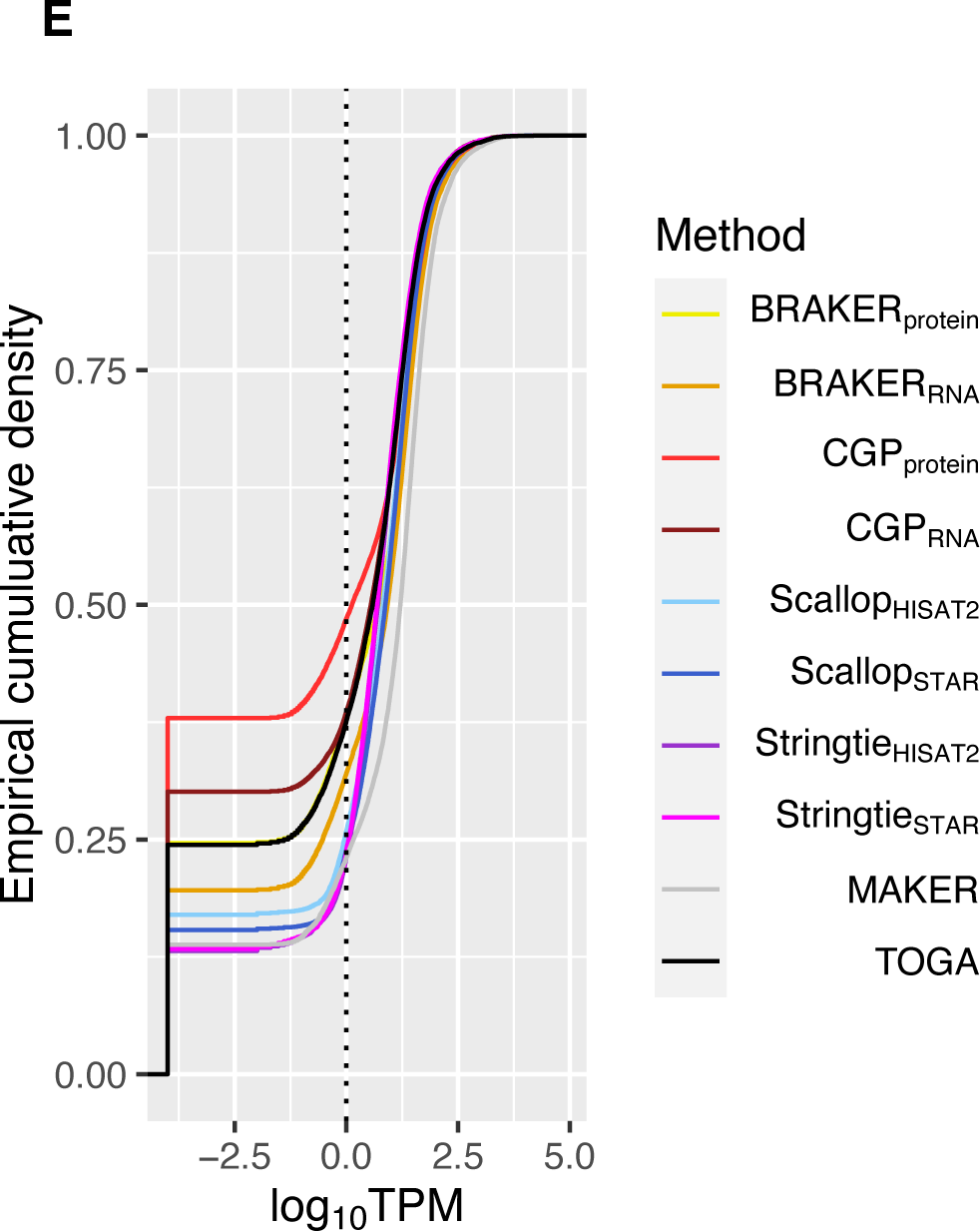
Empirical cumulative distributions of TPM for (A) *H. sapiens*, (B) *G. gallus*, (C) *A. thaliana*, (D) *Z. mays*, and (E) *D. melanogaster.* 0.0001 is added to all TPM values before log-transformation. Dashed line indicates TPM of 1.

**Figure S12.**
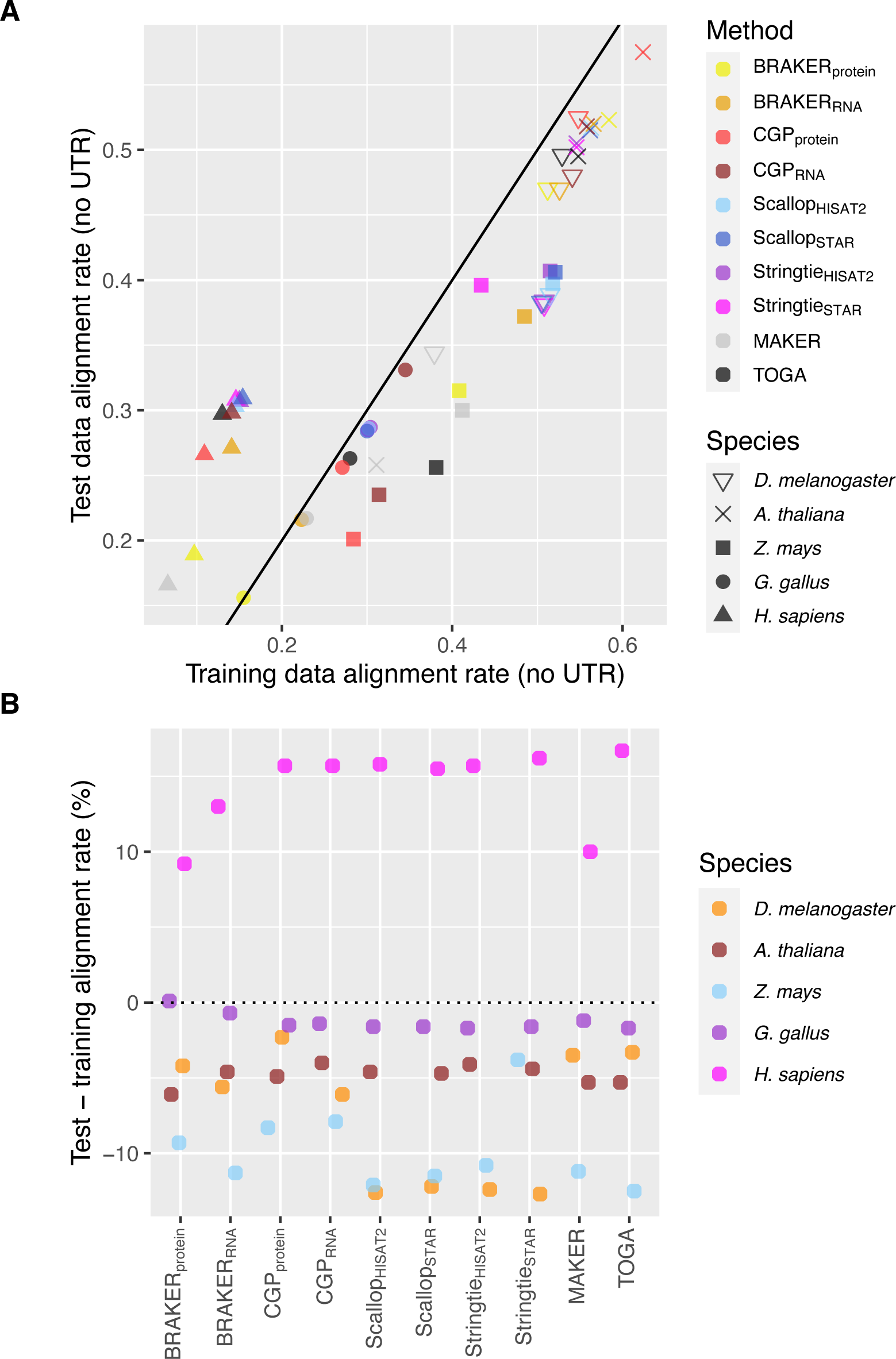
Comparison of RNA-seq alignment rates between training data used in annotations, and test data for the five reference species. (A) Pairwise plots of test again training rates, and (B) Rate differences by method a species.

**Figure S13.**
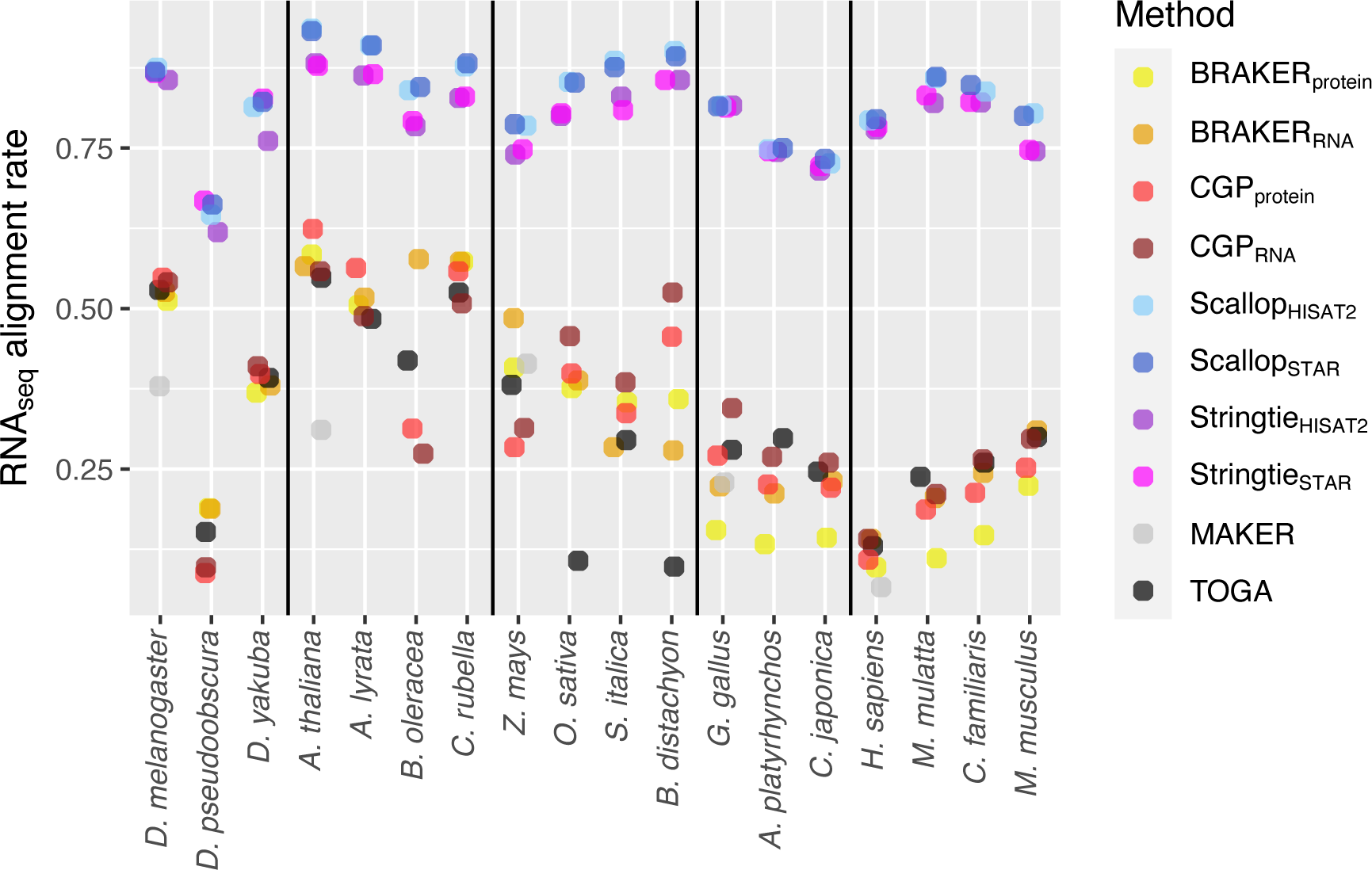
By method and species, RNA-seq read alignment rates to predicted transcripts, analogous to Figure 11, but for Stringtie and Scallop UTR intervals are included in the transcript predictions.

**Figure S14.**
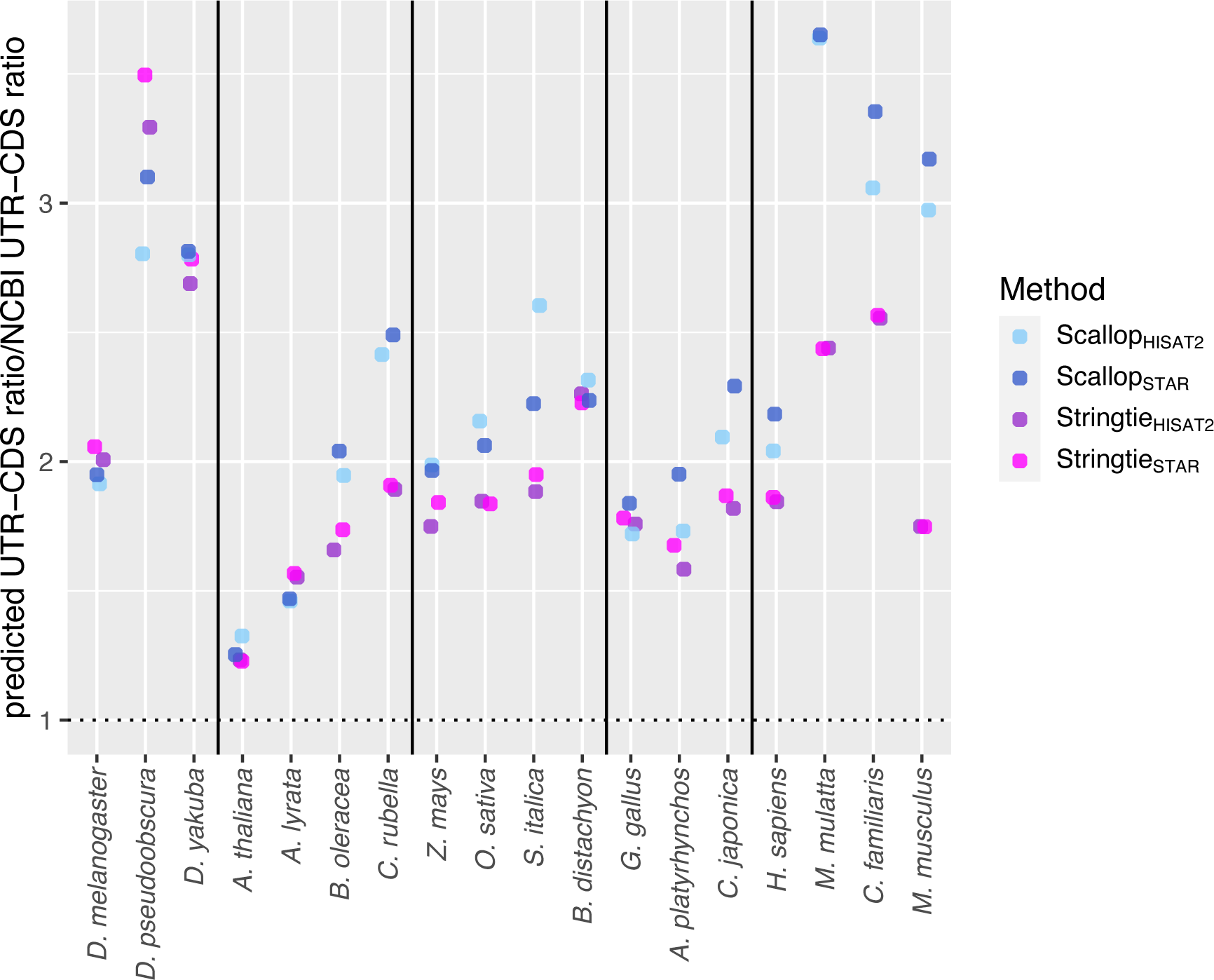
Ratios of the ratio of the median predicted UTR length/median predicted CDS length for RNA-seq assemblers over that for NCBI protein coding transcripts. No clear increase is observed for the most complete and curated annotations (*H. sapiens*, *D. melanogaster*, *G. gallus*, *Z. mays*, *A. thaliana*) relative to other genomes, indicating that the filtering out of cases of transcriptional readthrough (of the sort that NCBI will filter out for high quality annotations) does not explain the reduction in alignment rates for RNA-seq assemblers when UTRs are not included.

**Figure S15.**
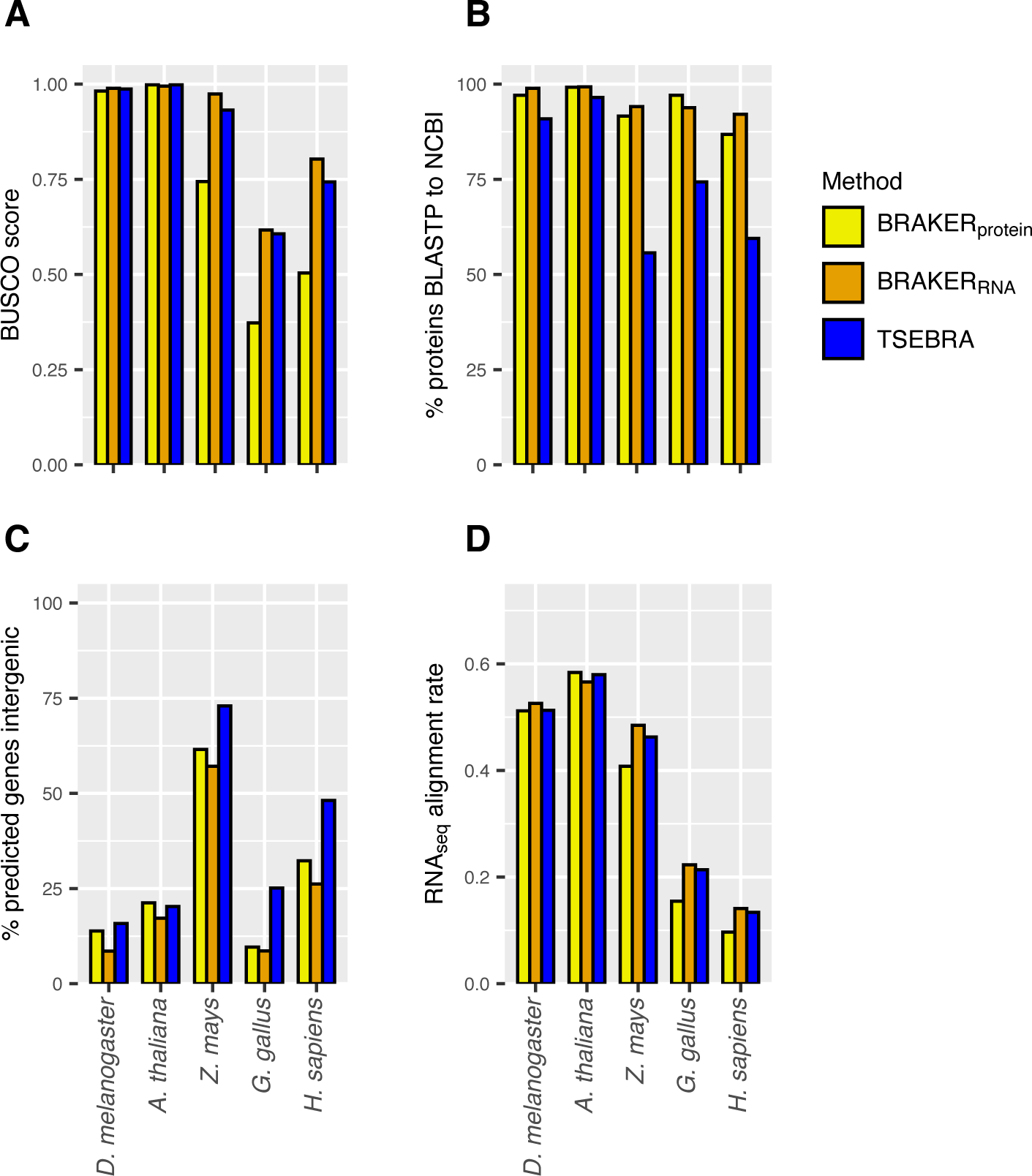
Comparison of TSEBRA integration to BRAKER_protein_ and BRAKER_RNA_ with respect to (A) BUSCO scores, (B) percentage of predicted proteins that have BLASTP hits to NCBI proteins for the species group, (C) the percentage of predicted genes that are intergenic relative to NCBI annotations, and (D) the RNA-seq alignment rates.

**Figure S16.**
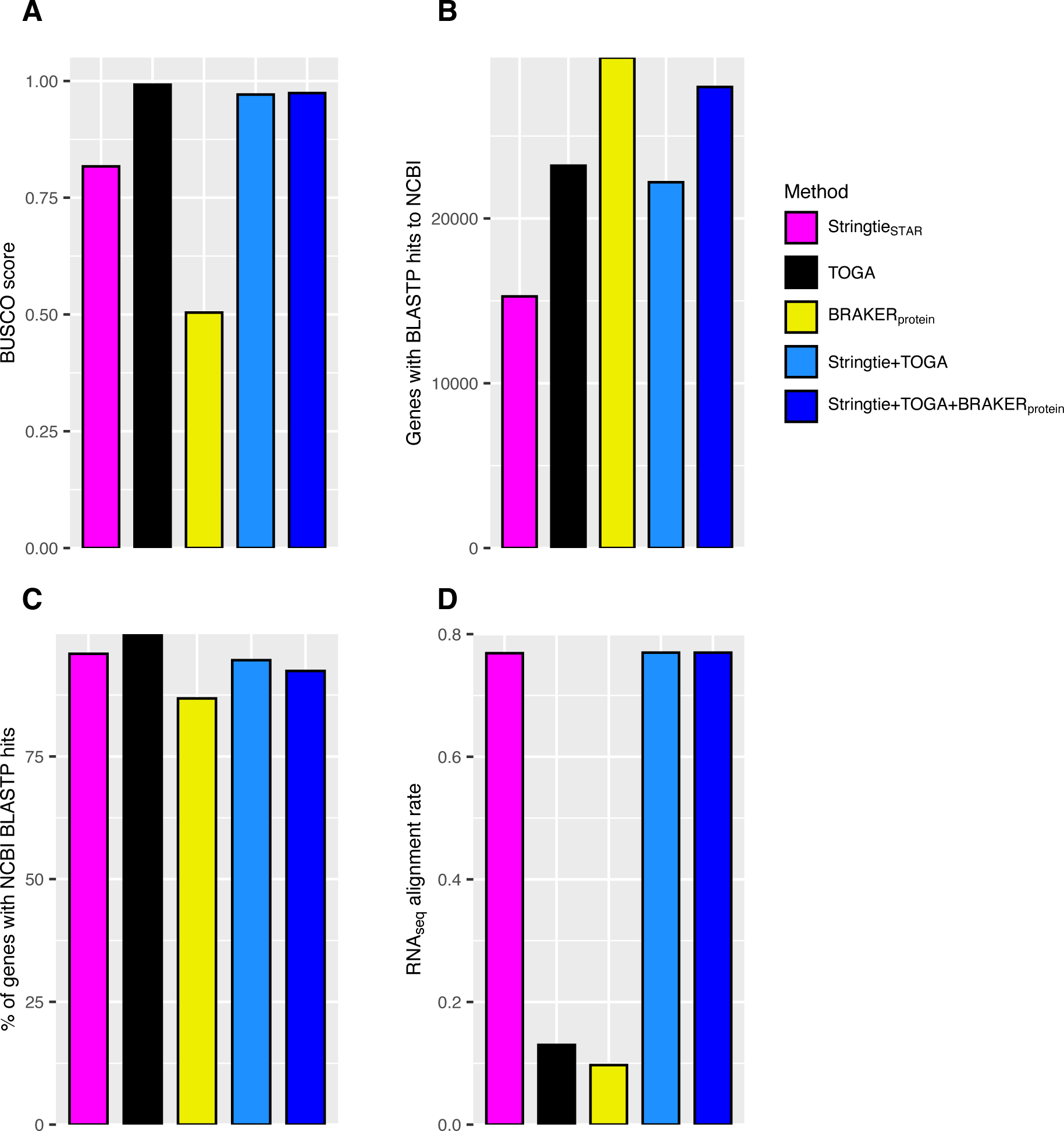
For *Homo sapiens*, Comparisons of (A) BUSCO scores, (B) the number of protein-coding genes with BLASTP hits to NCBI proteins from the mammal species group (C) the proportion of protein-coding genes with BLASTP hits, and (D) the median RNA-seq alignment rate across samples for individual methods and integrations of methods that start with Stringtie_STAR_, then add TOGA, and finally BRAKER_protein_. For each annotation that is added to the base annotation, integration involves adding genes that fall outside of the base annotation. For example, Stringtie+TOGA is built by adding TOGA genes that fall outside of Stringtie gene intervals.

**Figure S17.**
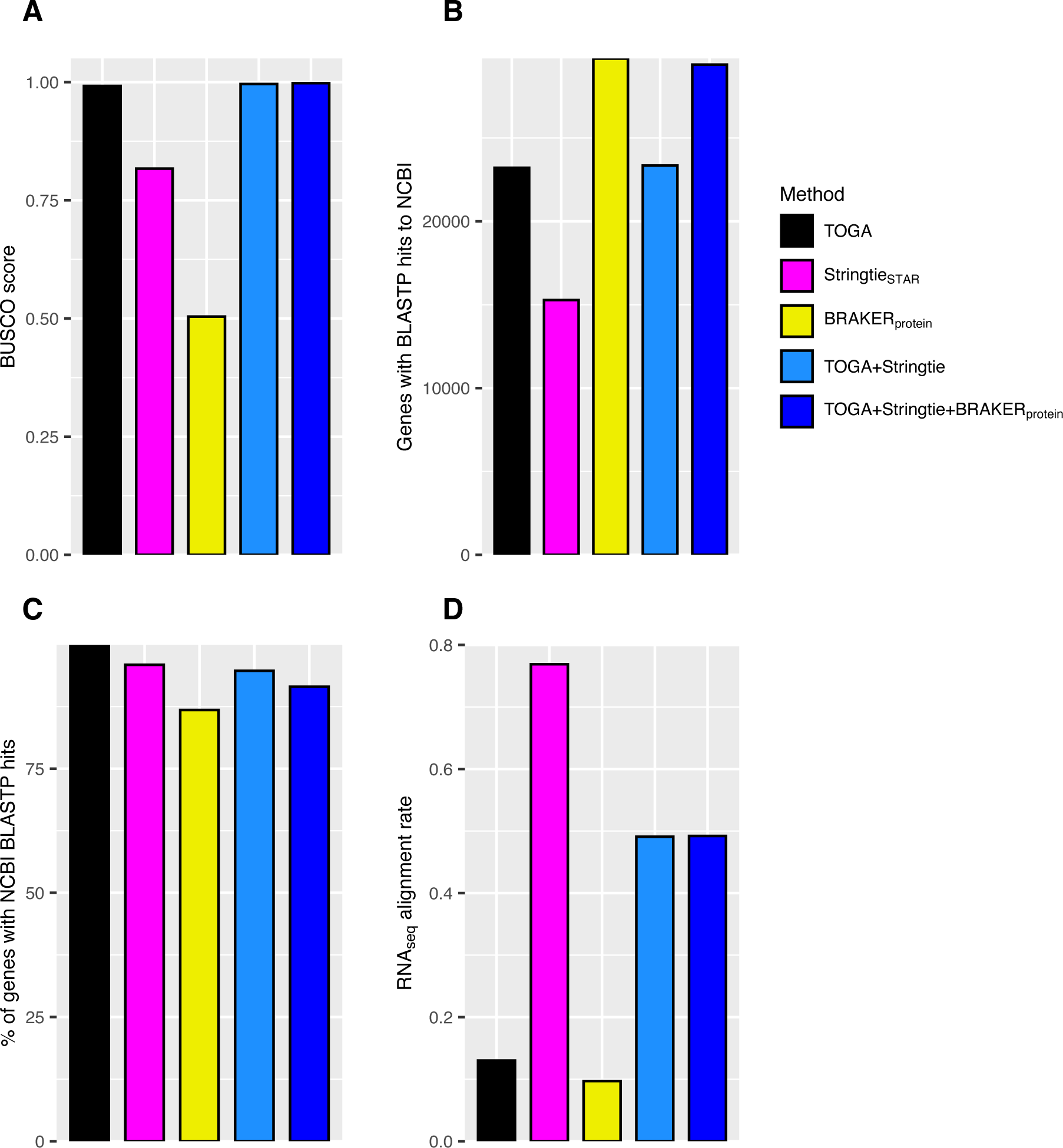
For *Homo sapiens*, Comparisons of (A) BUSCO scores, (B) the number of protein-coding genes with BLASTP hits to NCBI proteins from the mammal species group (C) the proportion of protein-coding genes with BLASTP hits, and (D) the median RNA-seq alignment rate across samples for individual methods and integrations of methods that start with TOGA, then add StringtieSTAR, and finally BRAKERprotein. For each annotation that is added to the base annotation, integration involves adding genes that fall outside of the base annotation. For example, TOGA+Stringtie is built by adding Stringtie genes that fall outside of TOGA gene intervals.

**Figure S18.**
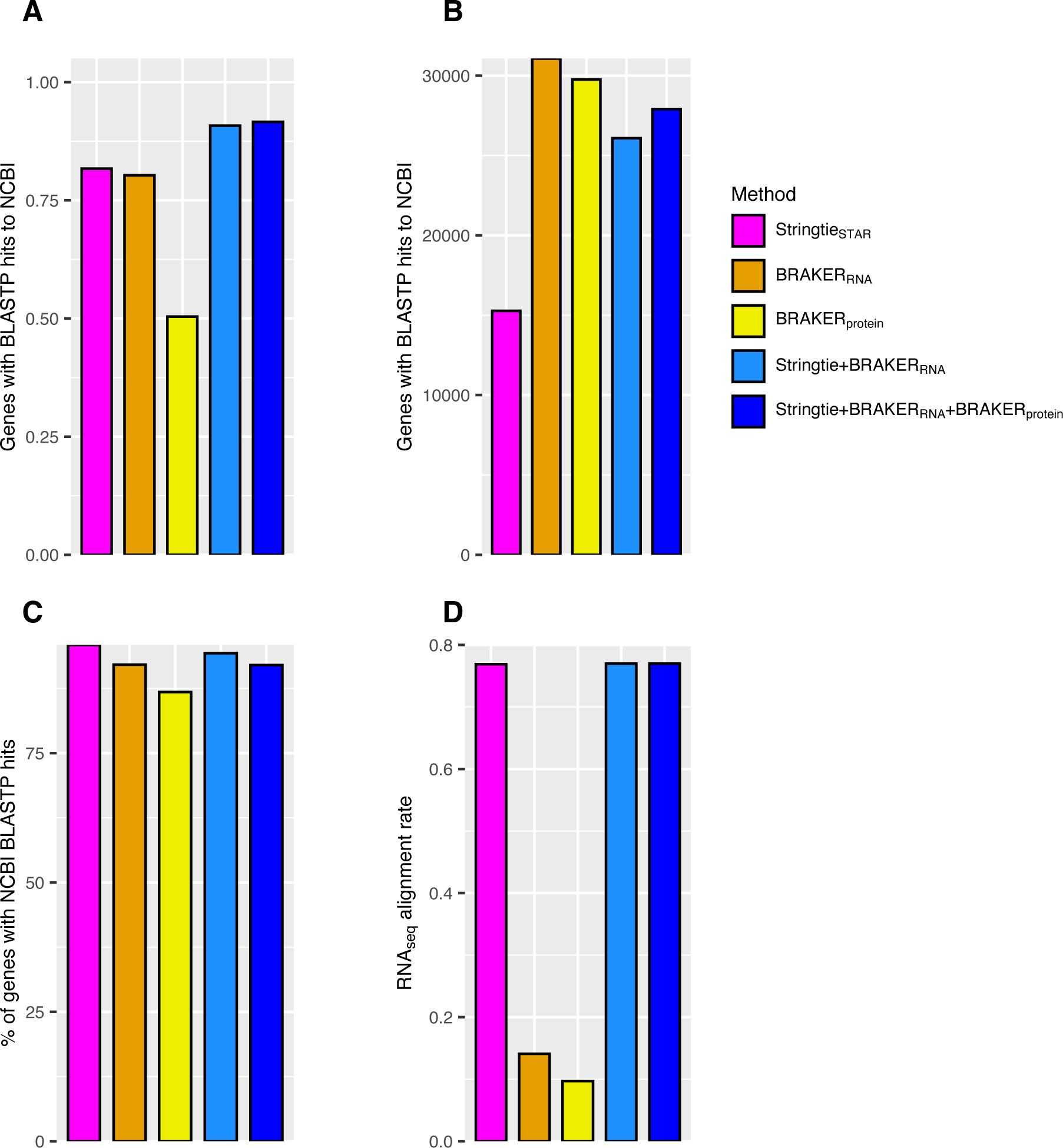
For *Homo sapiens*, Comparisons of (A) BUSCO scores, (B) the number of protein-coding genes with BLASTP hits to NCBI proteins from the mammal species group (C) the proportion of protein-coding genes with BLASTP hits, and (D) the median RNA-seq alignment rate across samples for individual methods and integrations of methods that use both RNA-seq and protein evidence but that don’t include annotation transfer with TOGA. For each annotation that is added to the base annotation, integration involves adding genes that fall outside of the base annotation. For example, TOGA+Stringtie is built by adding Stringtie genes that fall outside of TOGA gene intervals.

**Figure S19.**
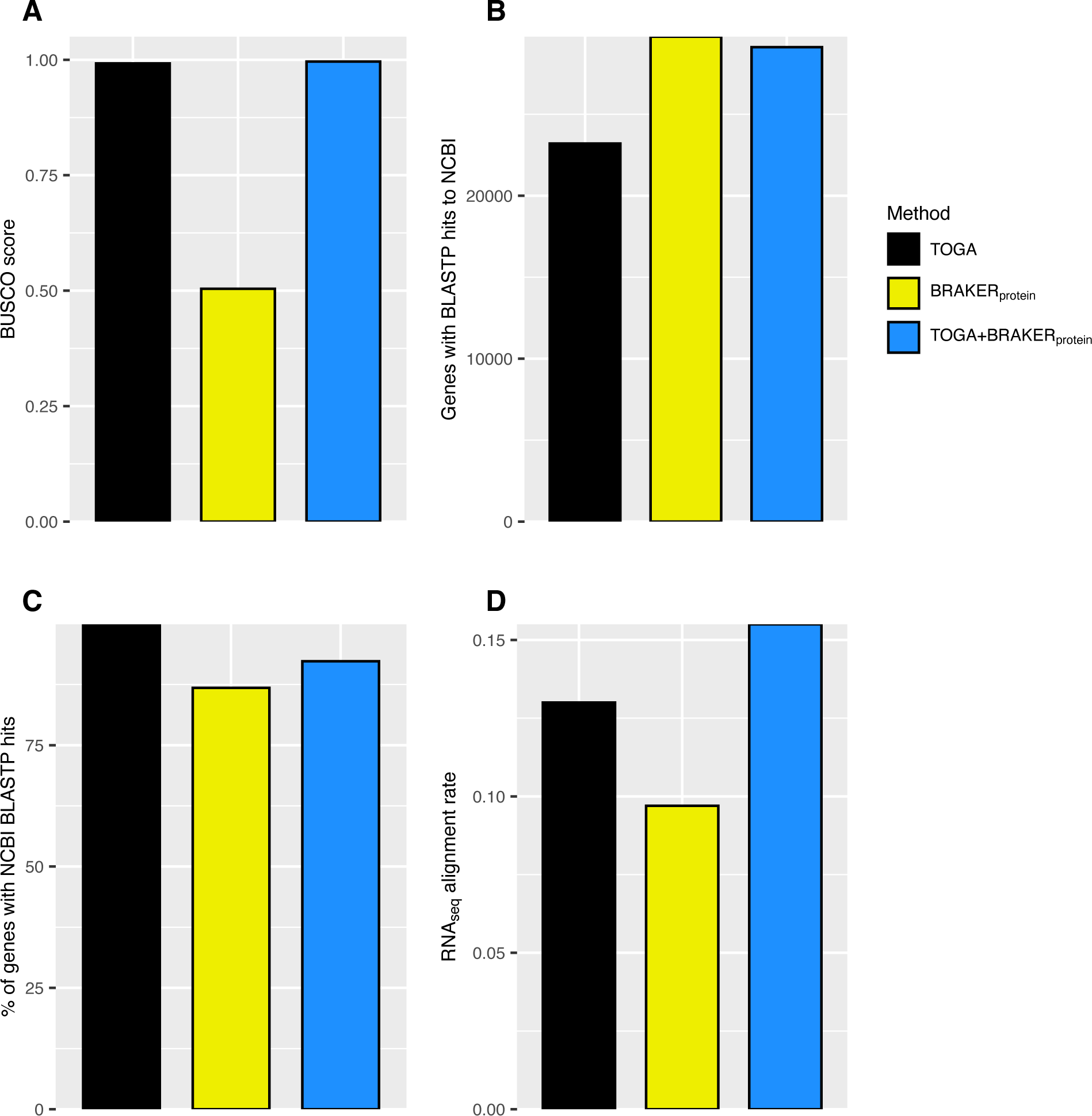
For *Homo sapiens*, Comparisons of (A) BUSCO scores, (B) the number of protein-coding genes with BLASTP hits to NCBI proteins from the mammal species group (C) the proportion of protein-coding genes with BLASTP hits, and (D) the median RNA-seq alignment rate across samples for individual methods and integrations of methods that do not include RNA-seq Integration of BRAKER_protein_ with TOGA involves adding BRAKER genes that fall outside of TOGA gene intervals.

